# Convergent molecular, cellular, and neural signatures of major depressive disorder

**DOI:** 10.1101/2020.02.10.942227

**Authors:** Kevin M Anderson, Meghan A Collins, Ru Kong, Kacey Fang, Jingwei Li, Tong He, Adam M Chekroud, B.T. Thomas Yeo, Avram J Holmes

## Abstract

Major depressive disorder emerges from the complex interactions of biological systems that span across genes and molecules through cells, circuits, networks, and behavior. Establishing how neurobiological processes coalesce to contribute to the onset and maintenance of depression requires a multi-scale approach, encompassing measures of brain structure and function as well as genetic and cell-specific genomic data. Here, we examined anatomical (cortical thickness) and functional (functional variability, global brain connectivity) correlates of depression and negative affect across three population-imaging datasets: UK Biobank, Genome Superstruct Project, and ENIGMA (combined N≥23,723). Integrative analyses incorporated measures of cortical gene expression, post-mortem patient transcriptional data, depression GWAS, and single-cell transcription. Neuroimaging correlates of depression and negative affect were consistent across the three independent datasets. Linking *ex-vivo* gene downregulation with *in-vivo* neuroimaging, we found that genomic correlates of depression-linked neuroimaging phenotypes tracked gene downregulation in post-mortem cortical tissue samples of patients with depression. Integrated analysis of single-cell and Allen Human Brain Atlas expression data implicated somatostatin interneurons and astrocytes as consistent cell associates of depression, through both *in-vivo* imaging and *ex-vivo* cortical gene dysregulation. Providing converging evidence for these observations, GWAS derived polygenic risk for depression was enriched for genes expressed in interneurons, but not glia. Underscoring the translational potential of multi-scale approaches, the genomic correlates of depression-linked brain function and structure were enriched for known and novel disorder relevant molecular pathways. These findings bridge across levels to connect specific genes, cell classes, and biological pathways to *in-vivo* imaging correlates of depression.

**Key Findings:**

1. Major depressive disorder and negative affect are associated with replicable profiles of cortical anatomy and function across independent population-level neuroimaging datasets (combined N≥23,723).
2. Somatostatin interneurons are consistent spatial transcriptional associates of *in-vivo* depression-linked imaging phenotypes.
3. Integrative single-cell gene expression analysis associate somatostatin interneurons and astrocytes with both *in-vivo* depression-linked imaging and *ex-vivo* gene downregulation in independent MDD cortical tissue samples.
4. Transcriptional correlates of *in-vivo* depression imaging phenotypes selectively capture gene downregulation in post-mortem tissue samples from patients with depression, but not other psychiatric disorders.
5. Indicating that some cell classes are preferentially sensitive to inherited disease liability, genome-wide risk for depression is enriched among interneurons, but not glia.
6. Gene associates of depression-linked anatomy and function identify specific neurotransmitter systems, molecular signaling pathways, and receptors, suggesting possible targets for pharmaceutical intervention.

## Introduction

Major depressive disorder (MDD) is a common and debilitating illness with a strong genetic basis (heritability, h^2^∼40%)^1^. Clinical depression emerges through complex interactions spanning multiple biological systems and levels of analysis^2^. The multi-scale nature of depression is evident in the presence of disorder relevant genetic loci^3^, as well as shifts in gene expression^4, 5^, cellular composition^6^, cortical anatomy^7^, and large-scale network function^8^. However, most research on the pathophysiology of depressive illness focuses on select features of brain biology, often in isolation. For instance, *in-vivo* neuroimaging studies link symptom profiles in patients to brain anatomy and network function^9, 10^, but are largely divorced from insights about underlying molecular and cellular mechanisms. By contrast, analyses of post-mortem tissue samples characterize illness related cellular and biological processes^4, 5, 11, 12^, but often focus on few regions and are limited by coarse diagnostic detail. To date there have been few opportunities to directly explore the depressive phenotype across levels of analysis — from genes and molecules through cells, circuits, networks, and behavior — simultaneously^13^.

*In-vivo* neuroimaging has identified depression related shifts in brain anatomy, metabolism, and function. For example, discoveries linking amygdala–medial prefrontal cortex (mPFC) circuitry to emotional^14^ and social processing^15^ led to the hypothesis that dysregulated interactions of cortical and subcortical systems precipitate the onset of depression^2, 16^. Further work has identified disrupted metabolism and altered grey matter volume in the mPFC of patients^17^, which may track illness chronicity^18^. However, as sample sizes have increased into the thousands it is now apparent that many early identified effects are likely more subtle than initially expected^7, 19^. As a consequence, the stability of depression relevant profiles of brain anatomy and function across populations remains unclear.

Complex clinical phenotypes like depression are tied to interactions throughout the functional connectome^8, 9, 20^. Supporting this perspective, biological subtypes and transdiagnostic features of depression may be revealed by considering the collective set of functional connections in the brain^9, 10^. Spatially diffuse correlates of depression across brain anatomy and function could arise from a host of biological changes in patient populations, ranging from demyelination^21^, altered neurotransmission^22^, and inflammation^23^, as well as changes in cell abundance or morphology^24^. Approaches that consider cross-level neurobiological shifts associated with depression would illuminate the biological bases of large-scale neuroimaging correlates of depressive illness. The recent emergence of whole-brain transcriptional atlases^25^ now permits more spatially comprehensive descriptions of the genomic correlates of depression, complementing targeted *ex-vivo* analyses of select cortical areas.

Post-mortem MDD patient data reveal abnormalities across cell classes, neurotransmitter systems, and molecular pathways^6, 24, 26^. For instance, diagnosis of depression is associated with reduced neuronal and glial cell size and abundance within prefrontal cortex^17, 27^ and subgenual aspects of mPFC^6, 24^. In particular, dysfunction of cortical somatostatin (SST) interneurons and astrocytes are hypothesized to play a preferential role in depression onset^11, 12, 28^. However, broad disruptions across molecular processes have been documented, including depression related dysregulation of pathways related to apoptotic stress and neuroinflammation^29, 30^, g-coupled protein receptors (GPCR) and cytokine activity^4^, ERK signaling and excitatory neuron activity^5^, as well as extensive alterations that encompass most major neurotransmitter signaling systems^26^. The breadth of observed neurochemical disruptions in depression makes parsimonious descriptions of the disorder difficult. Moreover, the degree of diagnostic specificity linking depression relevant patterns of brain anatomy and of function with any given cellular or molecular abnormality remain unclear.

In this study, we identify shared neurobiological signatures of depression that link anatomical, functional, cellular, transcriptional, and genetic levels of analysis. Across three imaging datasets (combined N≥23,723), we establish diffuse but replicable profiles of cortical brain anatomy and function associated with both major depressive disorder and individual differences in trait negative affect. Whole-cortex gene expression analyses revealed transcriptional associates of depression-linked *in-vivo* imaging phenotypes. Polygenic markers of SST interneurons reliably associated with imaging signatures of depression, both across datasets and imaging modalities. Indicating that normative patterns of gene expression reveal regional vulnerability of cortex in depression, the transcriptional signature of *in-vivo* depression phenotypes correlated with gene downregulation in independent MDD *ex-vivo* patient tissue samples. Suggesting a degree of diagnostic specificity, this effect was not present within post-mortem cortical tissue analyses of patients with schizophrenia, bipolar disorder, autism, or alcohol abuse disorder. Cell correlates of depression brain phenotypes were identified via joint analysis of single-cell expression data and spatial gene signatures of *in-vivo* imaging data. Across techniques, SST interneurons and astrocytes emerged as consistent cellular associates of depression, nominated through both *in-vivo* imaging and *ex-vivo* post-mortem gene downregulation. Suggesting that inherited risk for the disorder is preferentially conveyed via particular cellular pathways, polygenic risk for depression showed enrichment for genes expressed in interneurons, but not glia. Finally, we demonstrate that the transcriptional associates of depression-linked imaging markers may reveal clinically relevant information, including preferential importance of specific biological pathways and sensitivity to particular classes of neurotransmitters. Taken together, these data identify stable imaging correlates of depression, highlighting the role of somatostatin interneurons and astrocytes, and define a roadmap for future multi-scale neuroscience research on genomic bases of brain structure, function, and risk for depression.

## Results

### Neuroimaging correlates of depression and negative affect are consistent across populations

To characterize the nature and stability of cortical imaging markers of depression and trait negative affect, we analyzed structural and functional MRI data from three independent large-scale collection efforts: UK Biobank (UKB)^31^, ENIGMA^7^, and Brain Genome Superstruct Project (GSP)^32^. Three imaging measures were examined: cortical thickness, resting-state functional amplitude (RSFA), and global brain connectivity (GBC). In the UKB (n=15,150), lifetime history of depression was determined from questionnaires collected at the MRI scan visit (see *Methods*). The diagnostic validity of the UKB depression phenotype was thoroughly explored. Rates of single (5.22%, n=791), moderate (9.93%, n=1,505), and severe (4.17%, n=631) depression were significantly positively associated with trait neuroticism, depressive symptom severity, genetic risk for depression estimated through prior GWAS^3^, and rate of antidepressant prescription (Supplementary Figure 1). Replication ENIGMA data reflect meta-analytic results from Schmaal and colleagues^7^ showing shifts in cortical thickness in patients with depression (ns=1,206-1,302) relative to healthy comparison participants (ns=7,350-7,449). In the GSP sample (n=947), trait negative affect was measured in healthy young adults using five convergent self-report questionnaires associated with the experience of negative mood (See *Methods*).

We identified whole-brain shifts in brain structure and function that are consistent across neuroimaging samples, phenotypic measures (i.e. depression status and trait negative affect), and imaging modalities (Figure 1). Depression and trait negative affect were associated with distributed shifts in cortical thickness, RSFA, and GBC, measured across 200 symmetric ROIs from the parcellation of Schaefer and colleagues^33^. Across UKB, ENIGMA, and GSP imaging datasets, we observed consistent global patterns linked to the history of depression for thickness (*r_s_*=0.29, p=0.018, p*_spin_*=0.016), RSFA (*r_s_*=0.40, p=4.7e-9, p*_spin_*=5e-5), and GBC (*r_s_*=0.41, p=2.3e-9, p*_spin_*=1e-4; Figure 1a-c; Supplementary Figure 2). The significance of spatial correlations was tested using spin-based permutation tests^34, 35^. Consistent with the theorized core role for disrupted heteromodal association cortex functioning in psychiatric illness^13^, depression relevant shifts in RSFA and GBC were preferential to heteromodal relative to unimodal cortices. That is, depression-linked Cohen’s *d* for RSFA was greater in heteromodal (M=0.018±0.025(SD)) relative to unimodal cortex (−0.026±0.023; p*_perm_*=0.001). By contrast, GBC was lower in heteromodal (−0.003±0.021) relative to unimodal cortex (0.008±0.03, p*_perm_*=0.003; Supplementary Figure 2). This unimodal/heteromodal distinction replicated in the GSP sample for both RSFA (p*_perm_*=0.001) and GBC (p*_perm_*=0.001). Together, these data provide evidence for small, yet replicable shifts in cortical anatomy and function linked to both depression and trait levels of negative affect. A critical further question is the extent to which subtle depression-linked profiles of *in-vivo* brain anatomy and function reveal consistent genomic markers of illness risk. Across all genes, spatial Allen Human Brain Atlas (AHBA) transcriptional associates of the depression cortical phenotypes were highly consistent between cross-cohort measures of thickness (r=0.62, p<2.2e-16), RSFA (r=0.81, p<2.2e-16), and GBC (r=0.75, p<2.2e-16). Figure 1d shows the gene-wise Rank-Rank Hypergeometric Overlap for each modality.

**Figure 1:**
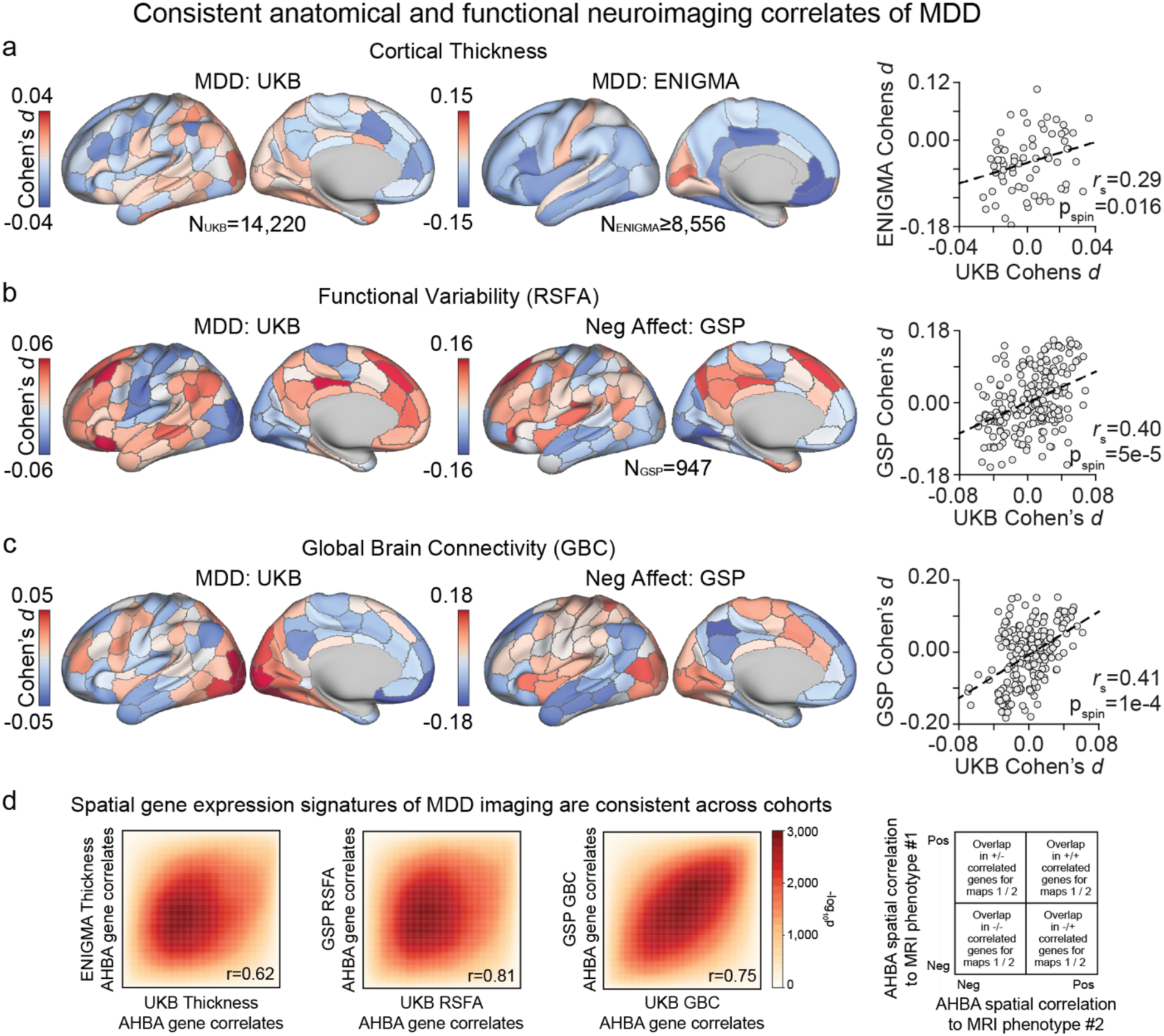
Imaging correlates of depression and negative affect are consistent across datasets. (a) Differences in cortical thickness between moderate/severe MDD and controls in the UKB and ENIGMA, and their spatial correlation (*r_s_*=0.29). Association of (b) functional variability and (c) global brain connectivity to moderate/severe MDD in the UKB, and negative affect in the GSP, and their spatial correlation (RSFA: *r_s_*=0.40, GBC: *r_s_*=0.41). (d) AHBA transcriptional correlates of MDD phenotypes are consistent across datasets: thickness (r=0.62), RSFA (r=0.81), and GBC (r=0.75). Heatmaps reflect -log10p calculated from Rank-Rank Hypergeometric Overlap tests.

### *In-vivo* depression imaging phenotypes track *ex-vivo* expression of somatostatin interneuron markers

Identifying molecular and cellular correlates of depression neuroimaging phenotypes would yield insight into the biological bases of the disorder and nominate targets for pharmacological intervention. For instance, brain areas that preferentially express genes related to a psychiatric disorder may be particularly vulnerable to illness progression^36, 37^. Reductions in somatostatin (SST) gene expression are a pronounced pathophysiological feature of depression^11, 38^, underscored by the preferential expression of SST markers in cortico-striatal reward circuitry and mPFC in donor tissue from healthy populations^36, 39^. SST expression is reduced within dorsolateral prefrontal cortex and subgenual mPFC in patients with depression^40, 41^, and experimental manipulation of SST neurotransmission in rodents modulates anxiolytic and antidepressant behaviors^42^ as well as socioaffective processing^43^. Depression linked alterations in the function of GABAergic cells, including SST, may influence signal-to-noise properties of cortex and global measures of connectivity^11, 38^, which could be reflected depression-related differences in RSFA and GBC. Thus, *in-vivo* imaging markers of depression may be most evident in cortical areas where SST related gene expression is greatest.

Next, we tested whether SST interneuron gene markers were spatially correlated to *in-vivo* patterns of brain anatomy and function associated with depression. Three highly selective gene markers of SST interneurons were analyzed (i.e. *SST*, *CORT*, *NPY*; Supplementary Figure 3 for validation of SST marker specificity). Critically, expression of SST markers should not be mistaken for a direct measure of cell abundance. Rather, relative expression likely reflects a combination of cell density and regional variability in cell transcription patterns. Cortical gene expression was measured with post-mortem AHBA data (see *Methods*). Figure 2a shows the normalized AHBA expression of *SST*, *CORT*, and *NPY* across 200 parcels (Supplemental Figure 4 for bi-hemispheric maps). Expression was also summarized across 68 desikan atlas parcels in order to match ENIGMA data.

**Figure 2:**
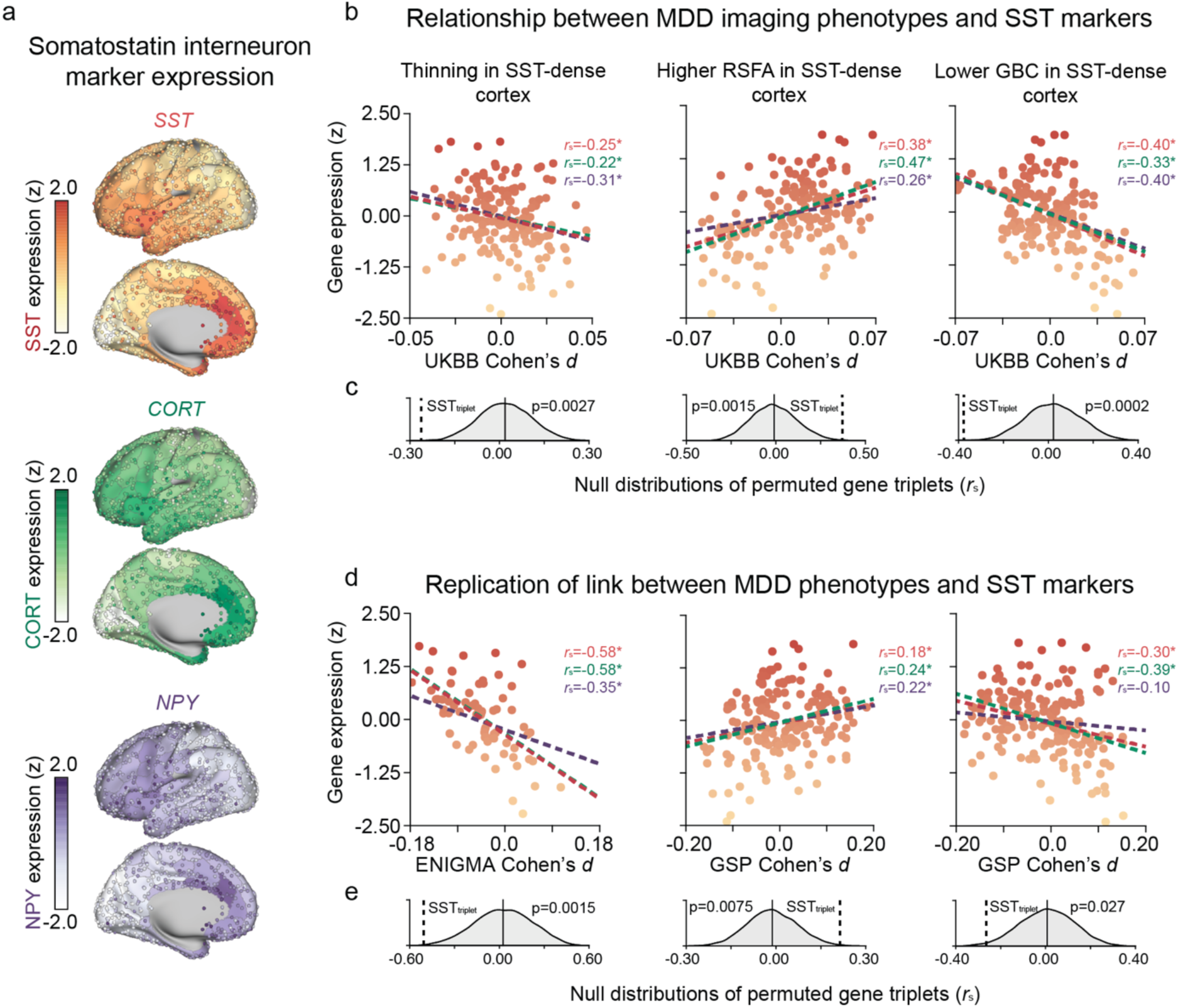
SST marker genes are spatially associated to *in-vivo* imaging correlates of depression. (a) Normalized AHBA cortical expression of three gene markers for somatostatin interneurons: somatostatin (*SST*), cortistatin (*CORT*), and neuropeptide Y (*NPY*). Each dot on the cortical surface represents expression in a single AHBA tissue sample, which is averaged across 200 bihemispheric cortical parcels. (b) SST marker expression is spatially correlated with depression-related shifts in cortical thickness (*r_avg_*=-0.25), RSFA (*r_avg_*=0.37), and GBC (*r_avg_*=-0.38) in UKB data. Circles in the dot plots are cortical parcels, colored by relative *SST* expression. Permutation analyses revealed that the strength of the spatial association was greater than what is expected by random selection of 10,000 triplets of brain-expressed genes. (e) SST marker spatial associations are consistent in replication data for cortical thickness (*r_avg_*=-0.51), RSFA (*r_avg_*=0.21), and GBC (*r_avg_*=-0.26). SST*_mark_*=average of *SST*, *NPY*, *CORT* spatial correlations.

Across all datasets and modalities, *ex-vivo* expression of SST gene markers spatially correlated to *in-vivo* depression-linked cortical phenotypes (Figure 2b-e). That is, *SST*, *CORT*, and *NPY* were expressed most in anterior cortical areas where depression-linked cortical thinning was greatest, an effect that is consistent in both UKB (*r_sst_*=-0.25, p*_spin_*=0.0004; *r_cort_*=-0.22, p*_spin_*=0.001; *r_npy_*=-0.31, p*_spin_*=1e-4) and ENIGMA data (*r_sst_*=-0.57, p*_spin_*=3e-4; *r_cort_*=-0.57, p*_spin_*=3e-4; *r_npy_*=-0.38, p*_spin_*=0.004). The strength of the association was also benchmarked against permuted gene triplets drawn from a pool of 17,448 brain expressed AHBA genes (two-sided p-value). Results were robust to alternative permutation strategies using sets of cell gene markers (Supplementary Figure 5). In terms of function, depression-linked increases in RSFA were greatest in areas with greater relative SST triplet marker expression, across both the UKB (*r_sst_*=0.38, p*_spin_*=1e-4; *r_cort_*=0.47, p*_spin_*=1e-4; *r_npy_*=0.26, p*_spin_*=1e-4) and GSP (*r_sst_*=0.18, p*_spin_*=0.007; *r_cort_*=0.24, p*_spin_*=8e-4; *r_npy_*=0.22, p*_spin_*=0.002) samples. For functional connectivity, SST triplet gene markers were significantly correlated with depression decreases in GBC across data from the UKB (*r_sst_*=-0.40, p*_spin_*=5e-5; *r_cort_*=-0.33, p*_spin_*<1e4; *r_npy_*=-0.40, p*_spin_*=5e-5) and the GSP (*r_sst_*=-0.30, p*_spin_*=5e-5; *r_cort_*=-0.39, p*_spin_*<1e-4; *r_npy_*=-0.01, p*_spin_*=0.09). For the first time, these data identify a replicable and parsimonious cell correlate of anatomical and functional neuroimaging signatures of depression, and further nominate SST interneurons as a potential target for biological intervention.

### Transcriptional associates of *in-vivo* depression imaging phenotypes capture patterns of *ex-vivo* gene downregulation in patients

An outstanding challenge is to link neuroimaging markers of psychiatric disorders to underlying molecular alterations. Towards this goal, we tested whether transcriptional associates of depression-linked cortical imaging phenotypes capture patterns of gene dysregulation in post-mortem samples of brain tissue from patients with depression. Normalized AHBA cortical gene expression was spatially correlated (spearman’s rho) to each of the six depression-linked structural and functional effect maps detailed above (Figure 3a). AHBA correlates were then averaged to obtain a 1×17,448 array reflecting gene-wise spatial association to depression cortical maps, where negative r-values indicate stronger association (e.g. increased intrinsic expression in areas of depression-linked cortical thinning). We also analyzed meta-analytic differential expression estimates from Gandal and colleagues^4^, reflecting the degree to which a gene is up- or down-regulated in post-mortem cortex of patients with depression, bipolar disorder (BD), autism spectrum disorder (ASD), alcohol abuse disorder (AAD), and schizophrenia (SCZ). This results in a gene-wise array for each disorder, where the *i*-th entry gives the degree to which the *i*-th gene is up or down-regulated in the diagnostic group (Figure 3b).

**Figure 3:**
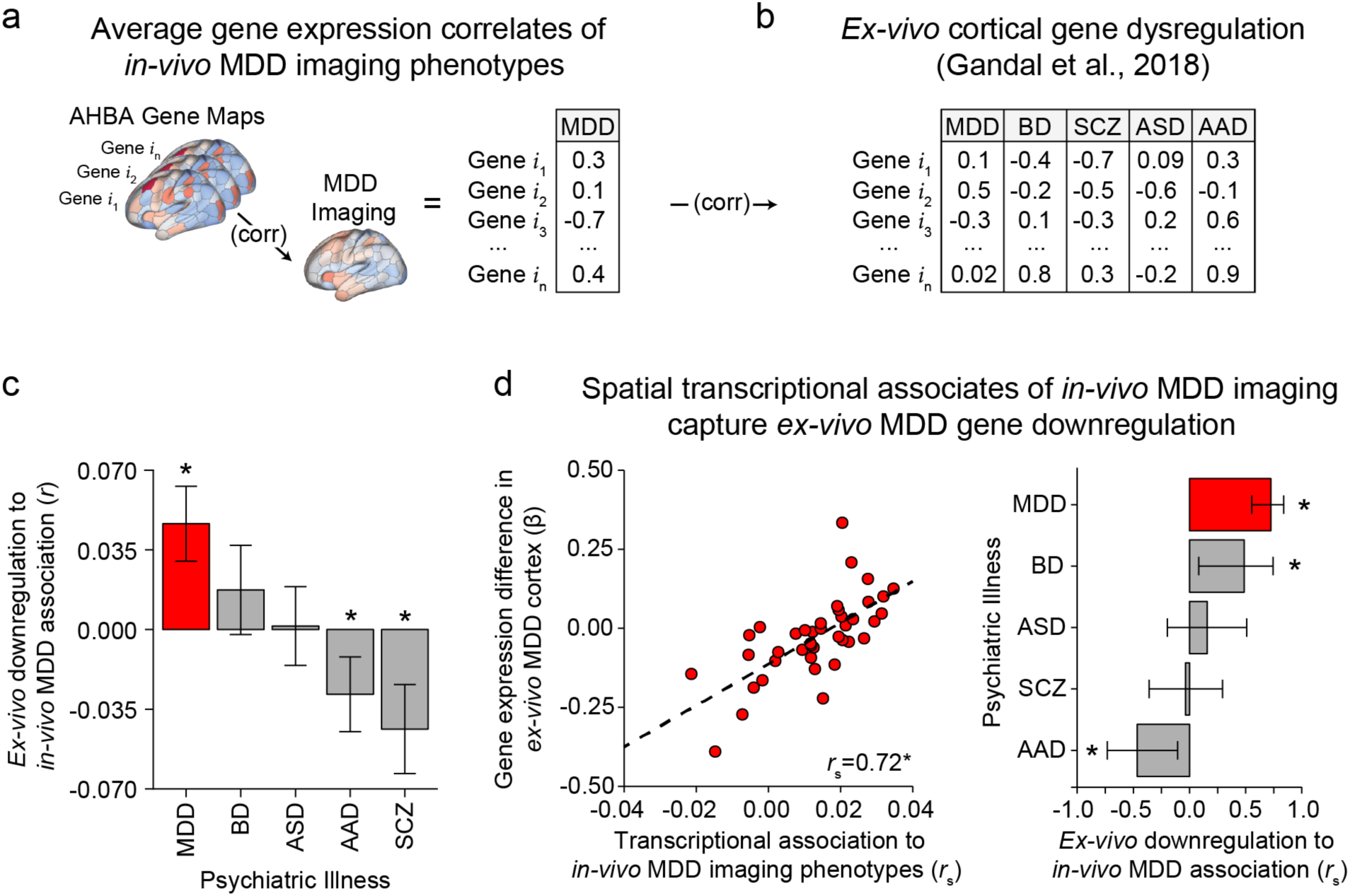
Selective association between *in-vivo* depression-linked imaging phenotypes and *ex-vivo* gene dysregulation in depression. (a) AHBA spatial gene expression was correlated to each the six depression-linked anatomical and functional neuroimaging map, then averaged. (b) Standardized case-control expression differences were calculated using post-mortem meta-analytic data from Gandal and colleagues^4^. (c) Average AHBA spatial correlation to depression maps was selectively correlated to post-mortem depression dysregulation (r=0.047, p=3.4e-8), but not to downregulation in other disorders. (d) Binned-analysis revealed a parallel relationship between gene downregulation in depression and AHBA correlates of *in-vivo* depression effects (*r_s_*=0.72, p=5.3e-7), which was also present for BD (*r_s_*=0.49, p=1.7e-3). MDD=Major Depressive Disorder; SCZ=Schizophrenia; ASD=Autism Spectrum Disorder; AAD=Alcohol Abuse Disorder. *p<0.05.

*Ex-vivo* cortical gene downregulation in depressed patients was significantly correlated to AHBA derived genomic associates of *in-vivo* depression cortical phenotypes (Figure 3a; r=0.047, p=3.4e-8). Suggesting a degree of diagnostic specificity, this positive relationship was selective to *ex-vivo* data from patients with depression and was not present in four comparison psychiatric disorders: schizophrenia (r=-0.044, p=1.2e-5), bipolar disorder (r=-0.017, p=0.082), alcohol abuse disorder (r=-0.028, p=0.0007), and autism spectrum disorder (r=0.0015, p=0.86; Figure 3c). Correlations for each disorder were calculated using all genes that were common across post-mortem and AHBA datasets. To explore the stability of this effect, we conducted a binned analysis relating *ex-vivo* dysregulation to AHBA imaging-genomic associates of depression (Figure 3d). For depression data, the 14,095 analyzable genes that were common across AHBA and Gandal et al.^4^ datasets were ranked by *ex-vivo* gene downregulation and divided into 40 gene bins. Average *ex-vivo* differential expression and average spatial association to *in-vivo* depression phenotypes was calculated for each bin, and then correlated. This approach revealed a significant correlation for depression data (*r_s_*=0.72, p=5.3e-7) that was highly stable across bin numbers, ranging from 10-40 in increments of 5 (range=0.69-0.92, M=0.78). The increased magnitude of this correlation (Figure 3c) is likely due to reductions in noise from binned-estimates relative to single-gene values. We also observed a significant effect for bipolar disorder (*r_s_*=0.49, p=0.002), but not other disorders. Together, these data indicate that areas marked by expression of genes that are downregulated in post-mortem patient tissue samples are more likely to show *in-vivo* illness-related shifts in brain structure and function (i.e., decreased thickness, decreased GBC, increased RSFA).

### Cell associates of *in-vivo* imaging phenotypes

The pathophysiology of depression is complex, emerging through interactions across multiple biological pathways and cell types^26^. Here, we incorporate single-cell expression data to explore the cell-type associates of depression-linked imaging phenotypes. A polygenic approach was adopted, since not all cell types express highly specific markers of their identity (e.g. *SST* in somatostatin interneurons). Cortical single-nucleus Droplet-based Sequencing (sn-Dropseq) data from Lake and colleagues^44^ were analyzed to identify positively differentially expressed genes across 16 transcriptionally defined cell classes (See *Methods*). This analysis resulted in 16 sets of gene cell markers (corrected q<0.05; See *Supplementary Data*). AHBA expression data were used to define spatial transcriptional associates of anatomical and functional neuroimaging correlates of depression (Figure 1). For each cell class, Fast-preranked Gene Set Enrichment Analyses (FGSEA) tested whether cell-specific genes were significantly more spatially correlated to anatomical and functional imaging markers of depression, normalized for the number of genes in a given cell-type set (i.e. Normalized Enrichment Score, NES)^45^.

Across all six imaging modalities and datasets, astrocytes, OPC, and Ex8 excitatory neurons (*CBLN2*+*POSTN*+) were significantly enriched for genes related to the derived depression neuroimaging phenotypes (Figure 4a). In line with *a priori* hypothesis driven analyses in Figure 2, SST interneurons were also positively enriched across 5 of the 6 imaging modalities. Astrocyte specific genes showed the strongest spatial association to depression neuroimaging effects (Figure 4b) and were expressed most within mPFC, anterior temporal lobes, and insular cortex (Figure 4c). The pattern of cell enrichment revealed by FGSEA was stable when compared to an alternative method, where the cortical expression of cell-specific genes was averaged using AHBA data (Figure 4c). In this approach, averaged cell-expression maps were then correlated to each depression imaging map. The cell-wise correspondence between this method and FGSEA was (*r*s=0.92, p<2.2e-16). We further established the stability of these results using polygenic cell deconvolution (Supplementary Figure 7)^46^. Deconvolution derived imputed distributions of SST interneurons and astrocytes were the two most spatially correlated cell types across all six modalities (rsst=-0.26, rast=-0.18).

**Figure 4:**
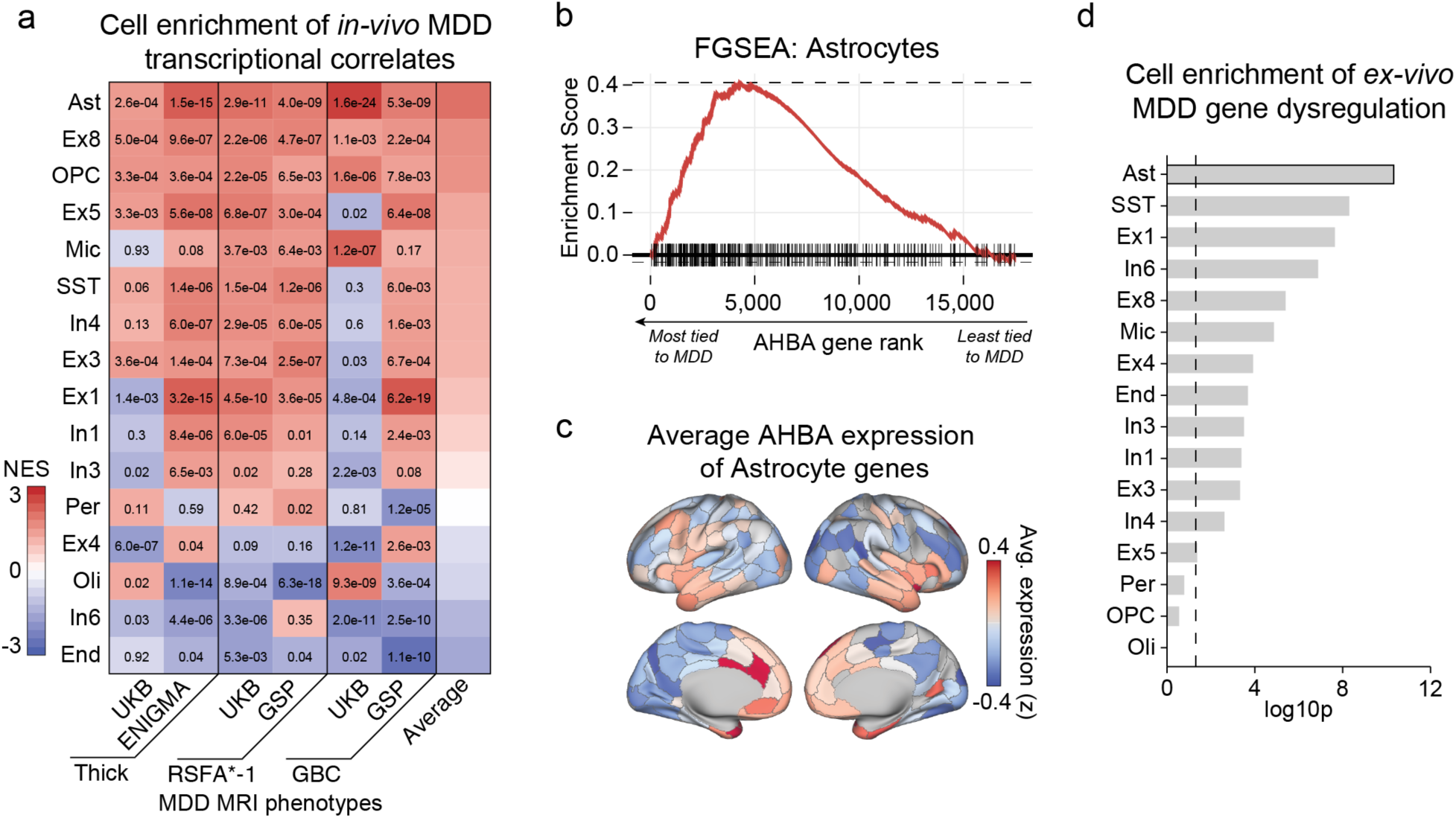
Integrative single-cell analyses implicate excitatory neurons, SST interneurons, and astrocytes. (a) AHBA genes were rank-ordered by spatial correlation to each depression MRI phenotype (e.g. UKB thickness, GSP RSFA). FGSEA identified astrocytes, OPC, and Ex8 (CBLN2+POSTN+) neurons as enriched across all modalities. RSFA gene correlates were multiplied by −1 to match the direction of thickness and GBC effects. Warm colors indicate positive enrichment, numbers reflect corrected p-values. (b) FGSEA enrichment plot showing that astrocyte marker genes tend to be spatially correlated to *in-vivo* depression maps. Each black line on the x-axis is the position of an astrocyte specific gene (c) Average AHBA expression of astrocyte marker genes, which was significantly spatially correlated to each depression imaging map (r_avg_=-0.20). (d) FGSEA analysis of genes downregulated in *ex-vivo* tissue samples from the cortex of patients revealed broad enrichment across cell classes, that was most pronounced in astrocytes and SST interneurons. NES=Normalized Enrichment Score.

The above results identify cell correlates of depression relevant neuroimaging phenotypes. However, any given cell class might be unchanged in depression or exhibit patterns of gene dysregulation. To address this, we conducted parallel FGSEA analyses using separate post-mortem data from Gandal and colleagues^4^ to identify cells enriched for gene downregulation in *ex-vivo* cortical tissue samples from patients with depression (Figure 4d). The NES values were above zero for most cells, indicating a broad pattern of depression linked downregulation amongst nearly all cell markers. The degree of observed cell enrichment was greatest for SST interneurons (NES=1.97, p=2.5e-09) and astrocytes (NES=1.97, p=8.2e-11). However, reduced astrocyte transcription was not a global feature of psychiatric illness, such that astrocyte markers showed significantly increased expression in SCZ (NES=-3.34), BP (NES=-3.10), and ASD (NES=-2.09) *ex-vivo* data. Taken together, these findings reveal patterns of reduced cell-specific gene expression in cortex of MDD patients that are greatest for SST interneurons and astrocytes.

Cell related abnormalities in MDD may reflect inherited genetic risk among cell-preferential pathways, or arise through environmental or second-order effects. Using GWAS data from Wray and colleagues^3^, we examined whether polygenic risk for depression is enriched among cell-preferential genes. Enrichment was measured with two methods, MAGMA gene-set property analysis^47^ and LDSC partitioned heritability^48^. Single cell expression data provided genomic signatures of 8 cell classes, measured from visual (V1C) and dorsal frontal (DFC) cortex^44^, as well replication data from temporal gyrus (MTG)^49^. Using LDSC, we observed significant enrichment of polygenic depression risk among interneuron specific genes in DFC (q=0.037) and MTG (q=0.46; Figure 5b). MAGMA revealed a similar pattern of enrichment for interneurons that was consistent across all three brain areas (V1C, q=1.4e-4; DFC, q=1.08e-3; MTG, q=3.4e-6). Excitatory neuron enrichment for depression GWAS signal was present with MAGMA, but not LDSC. We did not observe polygenic enrichment among any non-neuronal support cells, despite consistent associations of astrocytes to both *in-vivo* and *ex-vivo* depression phenotypes (Figure 4). Overall, our analyses suggest that areas with higher intrinsic expression of genes downregulated in depression were more likely to show illness-related shifts in both brain structure and function (i.e., decreased thickness, decreased GBC, increased RSFA).

**Figure 5:**
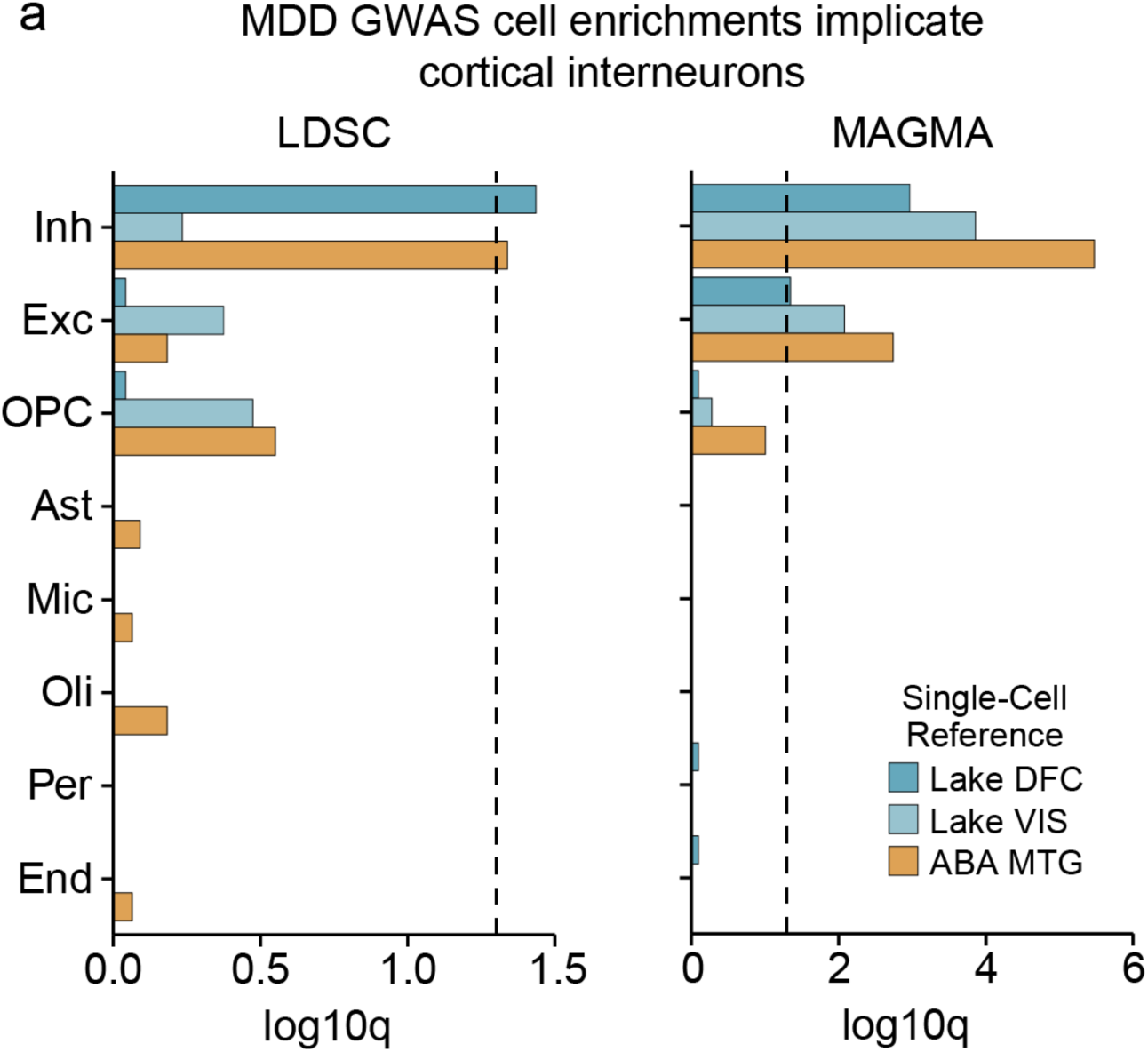
Genome-wide risk for depression is primarily enriched for inhibitory interneurons, but not glia. (a) Polygenic cell enrichment analyses were conducted across the eight superordinate cell categories across two methods, LDSC and MAGMA. Cell specific genes are defined using data from the MTG, DFC, and V1C. For LDSC, inhibitory interneuron markers show increased polygenic risk for depression^3^. For MAGMA, inhibitory and excitatory genes show enrichment for polygenic depression risk, but the effect is limited to differentially expressed genes defined from the MTG. Per=Pericytes; Oli=Oligodendrocytes; OPC=Oligodendrocyte Precursor Cells; End=Endothelial; Ast=Astrocytes; Inh=Inhibitory; Exc=Excitatory.

### Gene ontology of the transcriptional associates of depression

We next examined whether transcriptional associates of depression brain phenotypes capture clinically relevant information, such as sensitivity to a particular class of neurotransmitters, or increased importance of specific signaling pathways. Gene enrichment analyses were conducted using the top decile of genes correlated to neuroimaging markers of depression (n=1,745; Figure 6), revealing known and novel biological associates of depression. The top depression-linked gene decile possessed the greatest number of enrichment terms across molecular function, cellular component, and biological process ontological categories (Figure 6a). Further, genes related to “Depressive Disorder” and “Mental Depression” showed the strongest overlap with the top 10% of neuroimaging MDD gene correlates (Figure 6b). These effects indicate that coherent molecular processes capture MDD shifts in anatomy and function.

**Figure 6:**
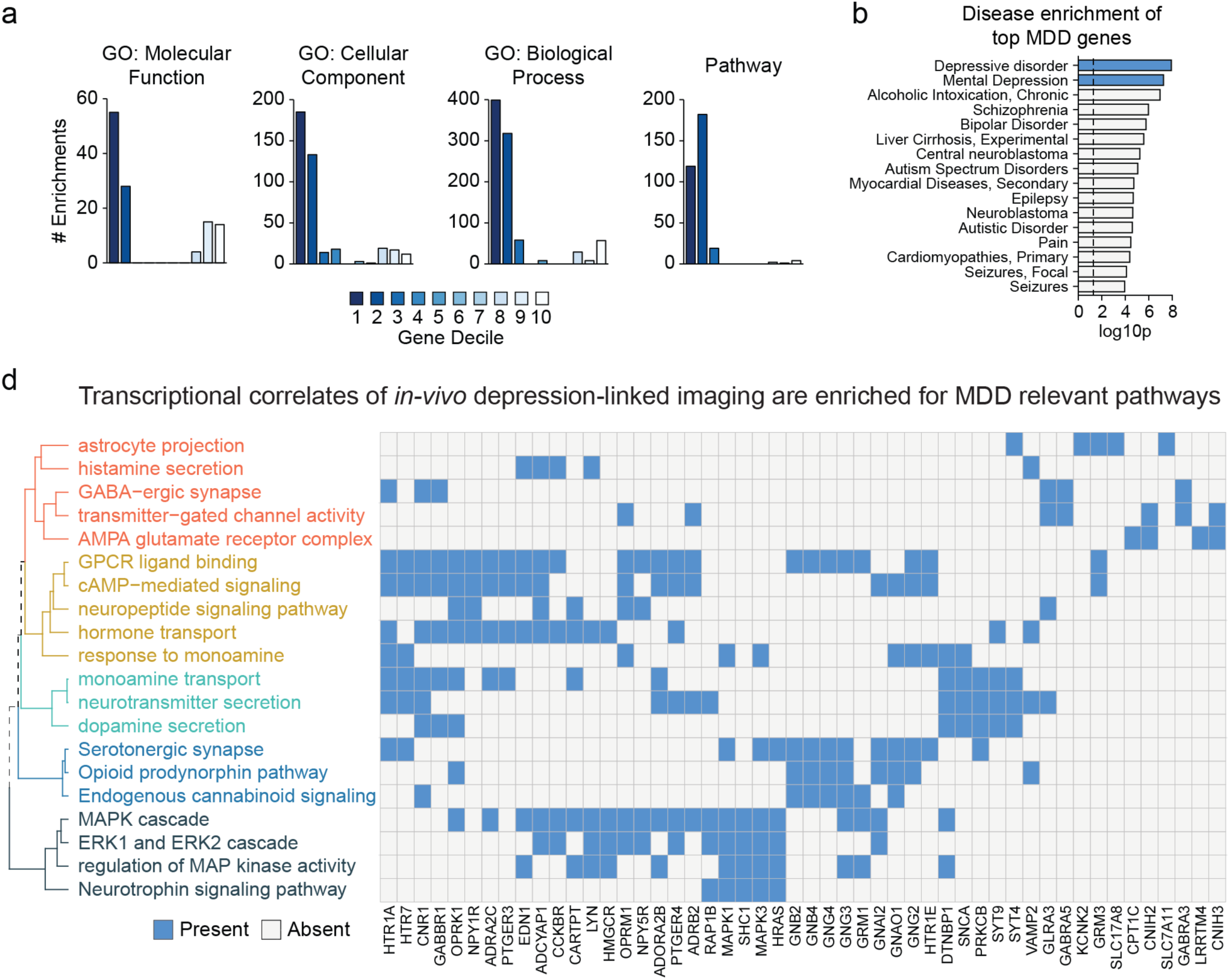
Transcriptional correlates of *in-vivo* depression-linked imaging phenotypes are enriched for depression relevant pathways. (a) Genes were rank-ordered by average spatial correlation to depression imaging maps, then split into deciles. The top gene deciles had the greatest number of enrichment terms across ontological categories. (b) The top gene decile was enriched for depression and other psychiatric disorders. (c) Subset of significant enrichment terms for the top decile of MDD imaging correlated genes. Hierarchical clustering is based on overlap of genes in each category. Blue indicates that the gene is included in a given enrichment term. Full enrichment terms are available in *Supplementary Data*.

Consistent with evidence of decreased glutamate and glutamine in the mPFC of patients with depression^50^, genes tied to glutamatergic receptors (GO:0008066, q=0.004) and secretion (GO:0014047, q=0.013) were significantly over-represented in the top decile. In line with prior reports of reduced cortical GABA in patients with depression^51^, we also observed enrichment terms for GABA receptor complex (GO:1902710, q=0.029) and GABAergic synaptic transmission (GO:0051932, q=0.041). Major monoamine neurotransmitter systems were also enriched, driven by genes tied to dopamine (GO:0014046; q=0.0098), histamine secretion (GO:0001821, q=0.036), and serotonergic synapses (Pathway 525336, q=0.017). Finally, our analyses identified terms related to g-protein coupled second messenger systems (GPCR; Pathway 1269544, q=0.00059) and downstream intracellular signaling pathways, including to cAMP-mediated signaling (GO:0019933, q=0.00033), ERK1 and ERK2 cascade (GO:0070371, q=0.043), MAPK cascade (GO:0000165, q=0.017), and non-canonical Wnt signaling pathways (GO:0035567, q=0.0044). These GPCR-activated intracellular cascades are important mediators of the neuromodulatory effects of neuropeptides^52^, which were also enriched in our data (GO:0007218, q=0.00021). Beyond SST, the neuropeptides substance P (*TAC1*), cholecystokinin (*CCK)*, cocaine-and-amphetamine related transcript (*CARTPT*), galanin (*GAL*), and receptors for mu and kappa opioids (*OPRM1*, *OPRK1*) in the top decile of MDD correlated genes. Critically, these results do not demonstrate that these systems are necessary altered in patient populations. Rather, they identify genes and pathways that are preferentially expressed within swaths of cortex implicated in depression.

These data are also consistent with gene networks implicated in depression from prior, independent, analyses of post-mortem cortical tissue. Specifically, there was strong spatial correlation between all depression imaging maps and the cortical expression of *DUSP6* (*r*avg=-0.365, 240^th^/17,448=0.003), which inhibits the ERK pathway and is a key hub gene that is downregulated within the mPFC of patients with depression^5^. Depression-linked neuroimaging effects were similarly correlated to *EMX1* expression, which is upregulated in the mPFC of patient populations. However, these spatial effects were in the opposite direction (*r_avg_*=-0.337, 17,034^th^/17,448=0.976), such that normative *EMX1* expression was lower in depression implicated areas of cortex (e.g. thinning of mPFC; Supplemental Figure 6). These findings suggest that expression differences in the cortical territories tied to depression may reflect divergence from normative patterns of area-specific expression, but more data is required to test this hypothesis.

## Discussion

The present analyses reveal converging biological signatures of depression that link neuroimaging, cellular, and molecular associates of the illness. Analyses of three population-imaging datasets identified replicable anatomical and functional cortical correlates of depression and negative affect. The observed neuroimaging markers of depression were spatially coupled to stable patterns of whole-cortex gene expression across both imaging modalities and data collections. Gene associates of *in-vivo* depression-linked cortical phenotypes were correlated with *ex-vivo* patterns of gene downregulation in cortex of patients with depression, but not samples from other comparison psychiatric disorders. In particular, gene markers of somatostatin interneurons and astrocytes were consistently spatially associated to *in-vivo* depression neuroimaging effects, and were downregulated within *ex-vivo* cortical samples from patients with depression. Indicating that some cell classes may be preferentially sensitive to inherited disease risk, cell enrichment analysis of depression GWAS data revealed increased polygenic burden among interneuron-specific genes, but not those of glia. Overall, we identify regionally variable imaging correlates of depression and present cross-modal data highlighting the particular role of somatostatin interneurons and astrocytes. Collectively, these results suggest potential biological targets for intervention and identify molecular pathways with exaggerated expression in depression implicated aspects of the cortex.

Our findings have important implications for understanding the cellular associates of structural, functional, and transcriptional alterations in depression. Prior neuroimaging research reveals subtle patterns of cortical thinning in mPFC, sgACC, and ventral temporal lobes that tracks illness severity^7, 17, 18^. In terms of function, depression is associated with reductions in global brain connectivity in mPFC and sgACC^53^, that extend to distributed aspects of multimodal association cortex^54^. The cellular bases of these alterations remain ambiguous, but may relate to GABAergic alterations^55^ or reduced size and density of neurons and glia^6, 27, 56^, particularly astrocytes^12^. Here, polygenic signatures of astrocytes were consistently associated to depression relevant shifts in both *in-vivo* imaging phenotypes and *ex-vivo* profiles of gene downregulation (Figure 4). Astrocytes influence synapse formation and elimination, as well as modulate neuronal communication, in part, through glutamate release and NMDAR receptor activation^57, 58^. Accordingly, depression related abnormalities in astrocytes may be involved in reduced glutamate levels in PFC and ACC of patients^12^. We did not find evidence for enriched depression polygenic risk among astrocyte specific genes (Figure 5), although speculative, this may suggest that observed cell alterations arise through environmental factors not directly tied to genetic risk. For instance, astrocytes are involved in neuroinflammatory signaling and are sensitive to cytokines, which are also implicated in depression etiology^12, 23^.

Considerable evidence indicates a preferential vulnerability of SST interneurons in depression and affective illness^11, 40^. Multi-scale studies in humans link SST related transcription to reward related cortico-striatal circuitry as well as regional variation in cortical function^36, 39^. Here, polygenic SST marker genes were significantly spatially correlated to all six depression neuroimaging phenotypes (Figure 2). That is, SST gene markers were expressed most in sgACC, mPFC, anterior insula and temporal lobes (Figure 2) corresponding to areas of depression-linked cortical thinning, increased amplitude, and decreased global connectivity (Figure 1). Highlighting the association of SST and astrocytes reported here, recent evidence indicates that astrocytes are particularly sensitive to SST interneuron activity, mediated in part by binding of somatostatin to astrocytic GABABR receptors^59, 60^.

In depression, SST related expression is consistently downregulated within the sgACC and amygdala of patients^28, 61^. Modulation of SST interneuron activity experimentally reduces depressive-like behavior in animal models of depression^42^, and is selectively tied to affective state discrimination in rodents^43^. Given evidence that SST expression is sensitive to BDNF and is cAMP dependent, depression related decreases in *SST* may reflect differences in neuronal activity rather than altered cell morphology or number^62, 63^. Spatial maps of *SST* marker expression shown here should not be conflated with a direct measure of SST cell density. However, rhesus macaque data indicate that the density of *CALB1* expressing interneurons (a subset of SST cells) and ratio of glia/neurons is highest within agranular limbic cortex relative to lateral PFC^64^, consistent with data in rodents^65^. Nonetheless, differences in relative expression of the neuropeptides like *SST* and *NPY* are likely functionally important, given their ability to influence neuronal and glial function^52, 59^. Increased relative expression of *SST* has further been documented among distributed whole-brain affective circuitry, including the nucleus accumbens, ventral tegmental area, mediodorsal thalamus, and anterior hippocampus^36, 66^. Future work should investigate whether alterations in SST cells in depression are consistent across distributed cortico-limbic circuitry.

Our results suggest that normative patterns of brain gene expression capture biologically meaningful information about depression related differences in cortex. These data converge with reports that spatial gene expression may reflect regional sensitivity to psychiatric illnesses^36, 67^, neurodevelopmental disorders^37^, and normative brain function and organization^39, 68, 69^. Here, we demonstrate that gene-wise association to depression imaging phenotypes correlated with gene downregulation in post-mortem patient tissue samples (Figure 3). Such findings support the emerging hypothesis of “transcriptional vulnerability”^37^, where brain regions with high baseline expression of disorder-linked genes are more likely to be affected over the course of an illness. The current results nominate specific receptors and signaling pathways linked to depression implicated brain areas, which may guide targets for biological interventions. Although a subset of our data include healthy young adults with varying levels of negative affect^32^, the current analyses largely reflect neuroimaging and transcriptional correlates at the depressive illness endpoints, and neurodevelopmental approaches are required to prospectively identify genomic bases of areal vulnerability.

Our *in-vivo* analyses identified consistent, yet subtle, anatomical and functional correlates of MDD and trait negative affect (Figure 1). The small effect sizes observed in UKB data could be due to the relatively older mean age of our sample or the use of self-report MDD symptoms (see Supplemental Figure 1 for validation of MDD phenotype). However, null hypothesis significance testing becomes problematic when analyzing very large samples, particularly in the UKB which has a target recruitment of 100,000 individuals^70^. For instance, the magnitude of linear effects linking brain features and behavior tend to be muted and generally require multivariate techniques to account for an appreciable amount of variance^31^, possibly reflecting the distributed nature of information processing in the brain^71^. Such a scenario echoes issues faced within the field of population genetics, where single genetic polymorphisms may have fleetingly small effects, but global or whole-genome analyses explain a considerable portion of trait variance^72^. Future work should investigate whether the pattern and magnitude of disease relevant neuroimaging effects vary by patient subpopulation or are differentially expressed across divergent symptom presentations^5^ or diagnosis constructs^73^, particularly given marked symptom heterogeneity seen in patients with depression.

A strength of the current analyses is our focus on global patterns of brain anatomy and function. For instance, our whole-cortex analyses revealed a surprising pattern of slightly increased visual cortex thickness in patients relative to controls (Figure 1), an effect that may be missed by small samples or hypothesis driven examinations of select brain areas. Of interest, this anatomical effect may, at least in part, explain patterns of increased functional connectivity in occipital cortex, reported both here through other collection efforts^73^. We also find that MDD and trait negative affect were associated with distributed functional changes that dissociate unimodal versus heteromodal cortex (Supplementary Figure 2). Global brain connectivity (GBC) was reduced in MDD across OFC, mPFC, and anterior temporal lobes, which contrasts increased GBC within visual cortex. These data support previous reports in smaller samples of reduced GBC in mPFC^53, 74^, but highlight the presence of broad spatially cohesive patterns of connectivity change. In this study, resting-state functional amplitude (RSFA) was increased in depression within heteromodal cortex, but was reduced in unimodal regions (Figure 1). Depression changes in RSFA and GBC were spatially anti-correlated to one another, supporting prior evidence that BOLD signal amplitude is predictive of within-subject change in functional connectivity^75^. In sum, these data identify spatially variable functional patterns across cortex in depression and negative affect, providing a neuroimaging foothold from which to interrogate underlying molecular and cellular associates.

## Future Directions

Future work should consider more refined characterizations of depressive illness to reflect the significant heterogeneity of the disorder, as well as to differentiate finer-grained phenotypes such as treatment resistance depression, early illness onset, or potential biotypes^9^. These cross-level data may also be used to explore psychiatric phenotypes that are not amenable to post-mortem analyses, such as premorbid illness risk or resilience. For instance, detailed neural signatures of depression risk will be available through large scale developmental neuroimaging collections, such as the Adolescent Brain Cognitive Development (ABCD) study^76^, which can be integrated with expanding atlases of developmental brain gene transcription^77^. Such efforts would shed light on disease etiology and are motivated by data from Schmaall and colleagues^7^ suggesting anatomical correlates of MDD are stronger in adolescence, relative to adulthood. The present results may also be used to explore treatment targeting strategies aimed at brain areas related in depressive disorder. Further, our analyses suggest that anatomical and functional correlates of depression extend beyond the limbic and affective brain systems typically implicated in depression (e.g. thicker visual cortex), which are not often targeted for post-mortem transcriptional analyses. Future work should aim for more regionally comprehensive genomic data in order to disentangle areal differences in depression brain correlates, such as anatomical and functional changes in visual cortex.

## Conclusion

In this study, we identify replicable anatomical and functional neuroimaging correlates of depression and trait negative affect, which serve as a foundation for integrative genomic analyses. Normative expression of polygenic SST interneuron markers in cortex were significantly spatially associated to depression correlates across all imaging modalities and datasets, in line with the hypothesized importance of this cell type in the disorder. Our data also suggest that the transcriptional associates of depression neuroimaging phenotypes capture global patterns of differential gene expression in depression, measured in *ex-vivo* patient cortical tissue. Incorporation of single-cell gene expression data showed that gene markers of SST interneurons and astrocytes were particularly strong spatial associates of depression imaging phenotypes, and were preferentially downregulated in post-mortem tissue samples from patient populations. Enrichment analyses of depression transcriptional associates identified multiple biological pathways, including neuropeptides, GPCR binding, and related intracellular MAPK, ERK, and cAMP signaling. Together, these data provide an integrative profile of the biology of depression that spans neuroscientific levels of analysis, connecting specific genes, cell classes, and molecular pathways to *in-vivo* imaging correlates of illness.

## Methods

### Major Depressive Phenotypes

#### UK Biobank

Lifetime history of depression was imputed using retrospective self-report questions collected during the imaging scan visit. The criteria for major depressive disorder followed the procedures of Smith and colleagues^78^. Individuals meeting criteria for single, moderate, or recurrent depression reported depressed mood (UKB Field 4598) or anhedonia (UKB Field 4631) for a week or more, with the longest period of anhedonia/depressed mood lasting two or more weeks (UKB Field 4609/5375). Single episode depression was characterized by endorsement of only one lifetime symptomatic period (UKB Field 4620/5386), and required individuals to have sought treatment through a general practitioner (GP; UKB Field: 2090) or psychiatrist (UKB Field 2100). Both recurrent and severe depressive diagnoses required two or more lifetime symptomatic episodes of anhedonia or depressed mood (UKB Field 4620/5386). Assignment of moderate depression required treatment seeking through GP (UKB Field 2090), but not a psychiatrist (UKB Field 2100), whereas assignment of recurrent depression required the opposite. For subsequent analyses, moderate and severe depression were reclassified into a binary indicator. Imaging data from individuals with a single lifetime episode depression were not analyzed.

Neuroticism scores were calculated by summing 12 self-report items from the UKB neuroticism inventory asking about trait levels of mood, irritability, worry, nerves, and guilt^79^. A subset of individuals completed an online behavioral battery assessing lifetime history of depressive symptoms. These online questions assessed history of sadness (UKB Field 20446), anhedonia (UKB Field 20441), sleep change (UKB Field 20532), feelings of worthlessness (UKB Field 20450), difficulty concentrating (UKB Field 20435), fatigue (UKB Field 20449), thoughts of death (UKB Field 20437), and weight change (UKB Field 20536). Self-reported anti-depressant usage was assessed in the same manner as Wray and colleagues^3^, which identified 41 types of pharmaceutical therapies for depression. Binary coding indicated whether individuals were taking at least one anti-depressant at the time of the scan visit (UKB Field 20003). Polygenic risk for depression was calculated for each UKB participant using GWAS data from Wray and colleagues^3^. Non-imputed genotype data were analyzed from 14,105 White/Non-Latino UKB subjects. Plink v1.9 was used to preprocess SNP data to remove variants with MAF < 0.05, hwe < 1e-6, variant missingness < 0.1, and sample missingness < 0.1^80^, resulting in 371,458 analyzable SNPs. Genetic-relatedness matrices were produced with GCTA and 20 genetic principle components were estimated for use as later covariates^81^. PRSice was used to calculate polygenic risk scores for each individual^82^ without application of a p-value threshold to filter GWAS variants. We note that 10.5% of MDD patients in the Wray and colleagues^3^ GWAS are from the UKB pilot data release, thus there is some overlap in our samples. Polygenic risk scores were compared between MDD groups, controlling for phenotypic and genetic covariates.

#### ENIGMA

The results from the ENIGMA meta-analytic study by Schmaal and colleagues^7^ were used for comparison against UKB anatomical effects. Cortical thickness changes in recurrent adult depression versus controls were analyzed, quantified in terms of Cohen’s *d* effect sizes across 68 Desikan atlas ROIs. Sample size varied by ROI: recurrent depression participants (N=1,206-1,302), control participants (N=7,350-7,450). Meta-analytic estimates controlled for sex, age, and scan center. Detailed information about patient and study demographics are published^7^.

#### Genome Superstruct Project

Trait negative affect was assessed in the same manner as Holmes and colleagues^19^. A single self-report measure was comprised of five scales related to history of negative emotion, including the NEO neuroticism scale^83^, the behavioral inhibition scale from the BIS/BAS^84^, reported mood disturbance assessed with the Profile of Mood State (POMS)^85^, the Spielberger State/Trait Anxiety Inventory (STAI)^86^, and measures of harm avoidance assessed with the Temperament and Character Inventory (TCI)^87^. Scores on each scale were z-transformed across individuals prior to averaging to generate a trait negative affect composite score.

### Neuroimaging Processing

#### UK Biobank

Structural and functional MRI data from the UKB were analyzed using an extended version of the standard UKB preprocessing pipeline (https://git.fmrib.ox.ac.uk/falmagro/UK_biobank_pipeline_v_1). Anatomical and functional data from 16,350 individuals were initially available for analysis after preprocessing. Data were collected on Siemens 3T Skyra and 32-channel receive head coil and were obtained from the UK Biobank (Project ID: 25163). T1-weighted structural scans were reconstructed from raw DICOMS (TR=2,000ms, TE=2.01ms, TI=880ms, flip angle=8 degrees, resolution=1 mm^3^). Minimally preprocessed resting-state fMRI scans were acquired using a multiband gradient echo EPI sequence (length=6 min, FOV=210mm, slices=64, TR=735ms, TE=39ms, resolution=2.4 mm^3^). Data were collected on multiple scanners across imaging centers in Cheadle and Newcastle, UK. Detailed imaging protocols are published^31^. Scans were not analyzed if they were marked as corrupted or “unusable” by UKB automated quality control tools.

A minimally modified version of the UKB processing pipeline (”UK_biobank_pipeline_v_1”) was implemented to allow for surface-based structural and functional imaging analyses. The T1 structural pipeline included gradient distortion correction (GDC), reduction of the image field of view (FOV), and combined linear and non-linear registration to a 1mm MNI152 “non-linear 6^th^ generation” group atlas. GDC, linear, and non-linear transforms to standard space were implemented via a single combined transformation step to reduce image distortions. A similar pipeline was implemented for T2 FLAIR images, with the additional use of BIANCA to estimate the volume of white-matter hyperintensities^88^. The UKB resting-state fMRI pipeline (“bb_functional_pipeline”) was applied, reflecting the following preprocessing steps implemented with MELODIC^89^: GDC unwarping, EPI fieldmap unwarping, MC-FLIRT motion correction^90^, grand-mean intensity normalization, and high-pass temporal filtering. Using the trained classifier from the UKB, ICA+FIX was applied in order to remove structured artefacts and motion confounds (24 parameters). *FSL feat* was not used to perform spatial censoring of EPI data based on SNR, which would lead to missing data in frontal and temporal poles when rest data are project to the cortical surface. Instead, a binarized T1-derived brain mask was applied to remove non-brain voxels. Cortical thickness was estimated using non-face-masked T1 images, given evidence that face-masking can alter Freesurfer derived estimates of brain morphometry^32^. Freesurfer v6.0.0 was used to derive estimates of individual cortical surfaces and produce vertex-wise estimates of cortical thickness^91^ We do not use the Freesurfer “FLAIRpial” option to use the T2 FLAIR image for pial surface delineation. Vertex-level data were summarized using the 200 parcel 17-network functional parcellation by Schaefer and colleagues^33^. To produce ROI-based estimates of Freesurfer cortical thickness, the group-space functional atlas was transformed to the native surface space of each individual and statistics were extracted using the Freesurfer “mris_anatomical_stats” utility.

Surface-based preprocessing of the resting data was then conducted using a previously published pipeline (https://github.com/ThomasYeoLab/CBIG)^92,93^. Processing steps included identification of outlier EPI frames with frame-wise head-motion greater than 0.3mm or DVARS greater than 75. Outliers included one frame before and two frames after a detected movement. Alignment of Freesurfer processed structural data and functional images was conducted with Freesurfer’s boundary-based registration software, using a high-contrast EPI volume as a functional reference. Segments of the BOLD run were marked for removal if they consisted of fewer than 5 contiguous frames. A run was not analyzed if more than 50% of frames were outliers. Nuisance regression was conducted using linear regression to remove effects of global signal, average white matter signal, average CSF signal, average ventricular signal, six head motion estimates (3 rotational, 3 translational), and all corresponding temporal derivatives. Frames that were marked as outliers were not considered during the nuisance regression step. Censored frames were then interpolated using least-squares spectral estimation prior to band-pass filtering (0.009 Hz *≤ f ≥* 0.08 Hz) and removal of linear trends. The resulting preprocessed volumetric data was then projected onto fsaverage6 surface space, smoothed with a 2mm full-width half-maximum kernel and finally downsampled into fsaverage5 vertex space. The same procedure was conducted on non-bandpassed volumetric data to generate RSFA estimates.

Parcellated surface-based estimates of RSFA (standard deviation of BOLD time-course) and functional connectivity were estimated using HCP workbench. RSFA values were z-transformed within individuals. A subject’s 200×200 functional connectivity estimates were Fisher-Z transformed. GBC was calculated as the average correlation of a given cortical parcel to all 199 other parcels. Quantitative quality control included row-wise deletion for missing thickness or resting-state estimates across the 200 cortical parcels. Individuals with average cortical thickness, RSFA, or GBC more than *±*3SD from the mean were removed. Similarly, we identified outliers at the level of individual parcels and individuals with thickness, RSFA, or GBC outliers in more than 5% of all cortical parcels were removed. Finally, individuals with outlier (*±*3SD) total brain size or white matter lesion volume were censored from further analyses. MDD case status was binarized to mark controls and individuals with moderate/severe lifetime history of depression. Individuals reporting only a single lifetime history of MDD were not analyzed. The final sample included (N=2,136) cases and (N=12,084) controls (age: 62.78±7.40; percent female=53.07).

#### Genome Superstruct Project

Neuroimaging and phenotypic data from the open-access Brain Genomics Superstruct Project (GSP) were obtained (https://dataverse.harvard.edu/dataverse/GSP)^32^. All individuals were healthy young adults of White/Non-Hispanic ancestry with no history of psychiatric illness. Imaging data were acquired across multiple Seimens Tim Trio scanners at Massachusetts General Hospital and Harvard University. Only data from individuals scanned using 12-channel phased array head coils were analyzed. Anatomical data were collected using a multi-echo T1w magnetization-prepared gradient-echo image (multi-echo MPRAGE; TR=2200ms; TI=1100ms; TE=1.54ms; FA=7°; 1.2mm^3^; FOV=230). T2w anatomical data were acquired in the same session using a turbo-spin-echo with high sampling efficiency (multi-echo MPRAGE^94^: TR=2800ms; TE=327ms; 1.2mm^3^; FOV=192), with a bandwidth matched to the T1w acquisition (652 Hz/pixel). Resting-state functional MRI data were acquired using a single 6-minute gradient echo planar imaging (EPI) sequence (TR=3000ms; TE=30ms; FA=85°; 3mm^3^, FOV=216; slices=47; interleaved foot-head acquisition=1, 3, …45, 47, 2, 4, …, 44, 46).

Anatomical data were processed using the Human Connectome Project version Freesurfer v5.3.0. T2w images were incorporated to improve accuracy of pial surface estimation (i.e. “T2pial” flag). Freesurfer anatomical segmentations were visually inspected to identify gross abnormalities. Surface-based preprocessing of resting-state BOLD data was conducted with the CBIG pipeline (https://github.com/ThomasYeoLab/CBIG). Preprocessing steps included removal of first 4 TRs, slice-time correction, and identification of head motion outliers (frame-wise displacement > 0.2, DVARS > 50). Above threshold frames were identified, along with one frame before and two frames after a detected movement. Frames were also marked for removal if not part of contiguous segment of five low-motion frames. A scan was not analyzed if more than 50% of frames were marked for removal. MCFLIRT was used to perform motion correction with spline interpolation. Freesurfer’s *bbregister* with FSL initialization was used to align T1w and EPI data, prior to generation of masks to mark brain voxels. Nuisance regression was performed with linear regression to remove effects of the global signal, mean ventricular signal, mean white matter signal, six motion parameters, and their temporal derivatives. Motion outlier frames were not considered when performing nuisance regression. Data were then band-pass filtered (0.009 Hz *≤ f ≥* 0.08 Hz). Volumetric data were also transformed to surface space without band-pass filtering for later calculation of RSFA.

Resulting preprocessed data were then projected onto fsaverage6 group space (2mm vertex spacing) using Freesurfer’s “vol2surf”, and then smoothed with a 6mm full-width half max kernel. Smoothed surface data were downsampled to fsaverage5 space (4mm vertex spacing). Non-bandpassed vertex-level data as summarized across 200 bihemispheric cortical parcels, then parcel-wise RSFA was calculated in the same manner as UKB data (i.e. standard deviation of BOLD time-course). Parcel-wise connectivity estimates were calculated using band-passed data. Individual 200×200 connectivity estimates were Fisher-Z transformed. GBC was calculated as the average correlation of a cortical parcel to all 199 other parcels.

Behavioral and neuroimaging quality control resulted in 947 individuals for subsequent analyses (age: 18-74, M=21.8±5.05; percent female: 54.38; Shipley IQ: 113.40*±*8.49). Individuals with missing GBC or RSFA data in any cortical parcel were not analyzed. We then identified individuals possessing GBC or RSFA outliers (*±*3SD) in greater than 5% of cortical parcels, or extreme outliers in global GBC or RSFA (*±*4SD). RSFA values were then z transformed within individuals.

### Regression linking depressive phenotypes to neuroimaging

Regression analyses were conducted at independently across the 200 bihemispheric cortical parcels. All quantitative variables were z-transformed. In the UKB, the effect of moderate/severe MDD history (0/1) on cortical thickness, RSFA, and GBC were estimated, covarying for sex, age, age^2^, age*sex, age^2^*sex, total brain size, volume of white matter hypointensities, self-reported ancestry, genetically estimated ancestry (White/Non-Hispanic or not), T1 inverse SNR, MRI run-wise average motion and inverse SNR, diastolic and systolic blood pressure, X/Y/Z position of brain in the scanner (center mass of brain mask), and UKB imaging acquisition center. Regression analyses in the GSP sample was conducted in a parallel fashion to predict RSFA and GBC from trait negative affect, controlling for age, sex, age, age^2^, age*sex, age^2^*sex, intracranial volume, height, weight, Shipley fluid intelligence, years of education, scanner bay, and scanner console version. Cohen’s *d* effect size estimates of depression status were calculated using the t-statistic and df from the regression:

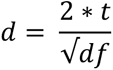

### Allen Human Brain Atlas

Publicly available human gene expression data from six postmortem donors (1 female), aged 24–57 years (42.5±13.38) were obtained from the Allen Institute. Data reflect the microarray normalization pipeline implemented in March 2013 (http://human.brain-map.org). Probes without Entrez IDs were removed and probe-wise noise for each donor was quantified as the number of above-threshold samples in cortex, divided by total cortical sample count. A probe-wise average was computed across all six donors, which was used to remove probes expressed in fewer than 20% of cortical samples. If more than one probe existed for a given gene, the one with the highest mean expression level was selected for further analysis, resulting in 17,448 brain-expressed genes. All analyses were conducted according to the guidelines of the Yale University Human Subjects Committee

Individual cortical tissue samples were mapped to each AHBA donor’s Freesurfer derived cortical surfaces, downloaded from Romero-Garcia and colleagues^95^. Native space midthickness surfaces were transformed into a common fsLR32k group space while maintaining the native cortical geometry of each individual donor. The native voxel coordinate of each tissue sample was mapped to the closest surface vertex using tools from the HCP workbench^96^. A cortical tissue sample was not analyzed if it was greater than 4mm from the nearest surface vertex, resulting in 1,683 analyzable cortical samples. Expression data was then averaged across 200 roughly symmetric surface ROIs from the 17-network functional parcellation of Schaefer and colleagues^33^. To allow for comparison to ENIGMA thickness data, gene expression was also summarized according to the 68 parcel Desikan atlas^97^. Even after normalization procedures employed by the Allen Institute to correct for batch effects, we observed residual differences in global expression intensity across cortical samples, possibly reflecting technical artifacts. Thus we perform within-sample z-transform normalization, similar to Burt and colleagues^98^, to reduce global expression differences across cortex. Microarray expression of each gene was then mean- and variance-normalized, revealing relative expression differences across cortex. Cortical data visualization was carried out using “*wb_view*” from the HCP workbench.

### Single-cell transcriptional enrichment analyses

We identify transcriptional markers of individual cell types using single-nucleus droplet-based sequencing (snDrop-seq) UMI counts for cells from visual (BA17) and dorsal frontal cortex (BA 6/9/10), obtained from GEO (GSE97942)^44^. UMI processing with Seurat was done separately for visual and frontal samples^99^. Initial filtering was conducted to ensure removal of genes expressed in fewer than 3 cells, as well as cells with fewer than 200 expressed genes. Expression values were normalized for each cell according to total expression values (i.e. “LogNormalize”), as well as covariates for sequencing platform and processing batch. Predefined superordinate cell categories from Lake and colleagues^44^ were used, which identified 16 cell classes that were present in both frontal and visual cortex. Differential expression in each cell type, relative to all others, was calculated using the Wilcoxon rank sum test in Seurat (i.e. “FindMarkers”). Seurat was used to conduct the same preprocessing steps on single-cell RNAseq data from the middle temporal gyrus (MTG) obtained from the Allen Institute^49^.

Cell enrichment analyses were conducted using Fast Gene Set Enrichment Analysis (FGSEA)^45^. Cell specific genes were identified based on significant positively differentially expressed in both frontal and visual cortex Lake data (fdr *≤* 0.05). Cell-wise FGSEA was then conducted for each neuroimaging modality (i.e. thickness, RSFA, GBC) and dataset (i.e. UKB, ENIGMA, GSP). Gene transcriptional associates of each imaging phenotype were identified using normalized AHBA expression data. That is, the cortical expression of each gene was spatially correlated to parcel-level depression neuroimaging effects (Cohen’s *d*). RSFA values were multiplied by −1 to align the direction of thickness and GBC effects. Ranked gene lists for FGSEA were in descending order based on spatial association to depression effects. Enrichment scores are the same that of GSEA^100^. We report normalized enrichment scores (NES) that account for the number of genes present in each cell marker group. Single-cell RNAseq MTG data were used as replication data^49^.

As technical replication, we also relate spatial averages of cell specific genes to each depression imaging phenotype. Simply, normalized spatial expression of AHBA cell specific genes from each cell class were averaged. The cell-wise expression maps were then spatially correlated to each anatomical and functional depression Cohen’s *d* effect map. Cell associations from the six imaging phenotypes were averaged, and then compared to the results of FGSEA. We also estimate spatial distributions of cell types by deconvolving bulk microarray AHBA data using CIBERSORTx (www.cibersortx.stanford.edu/)^46^. Gene symbols and entrez IDs were harmonized across AHBA and Lake snDrop-seq data using the NCBI gene alias dictionary (ftp://ftp.ncbi.nih.gov/gene/%20DATA/GENE_INFO/Mammalia/Homo_sapiens.gene_info.gz). Deconvolution analysis was performed using genes that were matched across single-cell and AHBA datasets. All data were transformed into non-log space, and single-cell gene signature matrices were produced separately for visual and frontal cortex snDrop-seq data. Each AHBA donor was deconvolved separately. Batch correction (“S-mode”) was performed to adjust for platform differences between UMI snDrop-seq and AHBA microarray data. Absolute imputed cell densities were mean- and variance-normalized separately for each AHBA donor, and then averaged across the Schaefer 17-network 200 parcel functional atlas and 68 node Desikan structural atlas. Individual cells were not analyzed if their imputed density was zero in more than 50% of cortical samples.

### AHBA spatial correlation to depression-linked neuroimaging

Normalized AHBA expression data, summarized by surface atlas parcels, was spatially correlated (spearman’s) to each of the six depression-linked neuroimaging phenotypes. We specifically investigate whether gene markers of somatostatin interneurons are spatially associated to anatomical and functional correlates of depression. The genes Cortistatin (*CORT*), Neuropeptide Y (*NPY*), and Somatostatin (*SST*) were used to index somatostatin interneuron transcription, given that they are highly and selectively expressed within SST interneurons. Given spatial autocorrelation among AHBA expression data, the significance of each expression-to-imaging correlation was assessed using spin-based permutation tests, which preserve the proximity-based correlation structure of expression maps^34^. We also perform multiple gene-based permutations to benchmark the strength of the association between SST gene markers (i.e. *CORT, NPY, SST*) and each depression-linked imaging phenotype. The first permutation randomly selects gene triplets (n=10,000 perms) from the pool of 17,445 brain-expressed genes (this exclude *CORT, NPY, SST)*. The second selects gene triplets (n=10,000 perms) from a select pool of 1,609 genes that were identified as significant cell type markers according to single-cell data from Lake and colleagues^44^ (excluding markers of SST interneurons). The third permutation strategy selects genes from the same pool of 1,609 marker genes, however each triplet is composed of genes that are significant markers for the same cell type (n=10,000 perms per cell type, excluding markers of SST interneurons). The results of each permutation strategy are presented in Supplementary Figure 5.

### Ex-vivo psychiatric patient differential expression

Meta-analytic estimates of differential expression from Gandal and colleagues^4^ were analyzed. Gene expression values were normalized prior to differential expression calculation with linear-mixed effects modeling in order to provide standardized beta coefficients, indicating the degree that a gene is up- or down-regulated for a given psychiatric population. Analyses included cortical expression data from patients with Major Depressive Disorder (MDD), Autism Spectrum Disorder (ASD), Bipolar Disorder (BD), Alcohol Abuse Disorder (AAD), and Schizophrenia (SCZ). Information about data preprocessing is published^4^, and sample information is available in *Supplementary Data*. Gene-wise patient differential expression (i.e. normalized beta) was then correlated to the gene-wise spatial correlation to *in-vivo* neuroimaging phenotypes.

### Single-cell MDD GWAS enrichment

We tested whether polygenic risk for depression, using the GWAS from Wray and colleagues^3^ was enriched among cell-specific genes. Enrichment was measured using MAGMA gene-set property analysis^47^ and LDSC partitioned heritability^101^. Cell-specificity of gene expression was measured with single-cell data from V1C, DFC, and MTG. Cell specific expression was quantified using the EWCE R package^102^. Genes were split into deciles for each cell, ordered from most specifically expressed to least. LDSC annotation files were created using the 1000 Genomes European Phase 3 release. Cell enrichment estimates were conditional on a baseline model (”1000G_EUR_Phase3_baseline”) of 53 genomic regions (e.g. enhancer, genic, etc.). Enrichment statistics were reported for the top gene decile for each dataset and cell type. MAGMA gene-property analyses followed those of Watanabe and colleagues^103^. We calculated averaged gene expression for each cell type and included overall gene-wise expression, collapsed across all cells, as a covariate. This approach also does not depend upon the creation of gene bins. For both methods, p-values were corrected for multiple comparisons (Benjamini-Hochberg) separately for each single-cell dataset (e.g. 8 tests for DFC snDrop-seq data).

### Gene Ontology Enrichment Analysis

ToppGene^104^ was used to identify biological enrichment terms across the MDD gene deciles. Genes were rank-ordered based on their average AHBA spatial correlation, collapsed across the six depression-linked neuroimaging maps, and then split into evenly sized gene deciles. The number of enrichment terms for each gene decile were then compared (Figure 6a), split across major ontological categories (e.g. Biological Process, Cell Component, etc.). We illustrate specific genes enriched among a circumscribed set of enrichment categories (Figure 6c). Similarity of each enrichment term was defined as the number of overlapping genes between two categories, relative to the number of total genes link to both enrichment categories. This similarity matrix was then hierarchically clustered to identify clusters of similar enrichment terms. Genes were selected for plotting if they were present among multiple enrichment categories.

### Code availability

Data and code used in this analysis are publicly available upon publication, unless restricted by data use agreement.

## SUPPLEMENTAL INFORMATION

**Supplemental Figure 1:**
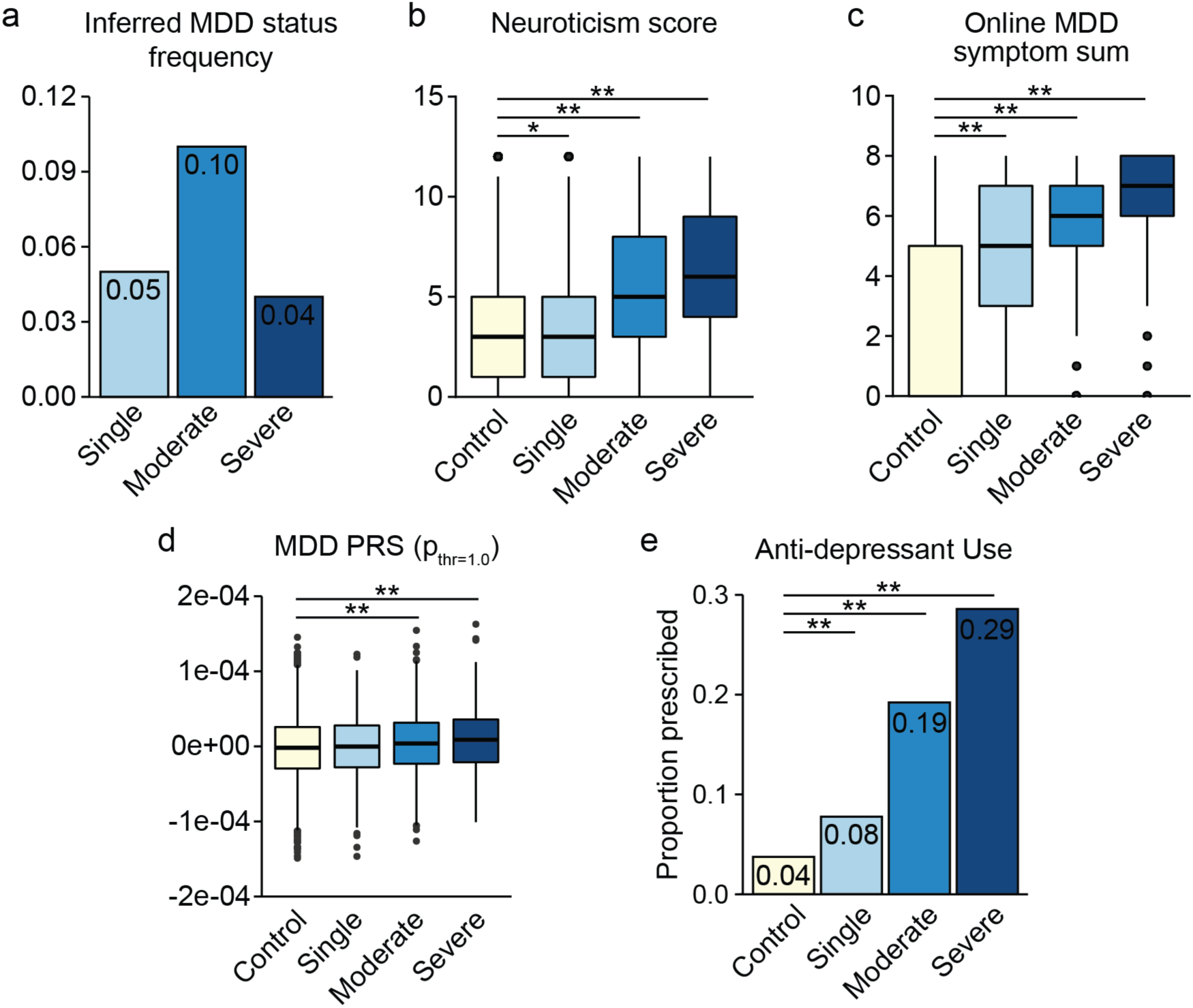
Construct validity of inferred depression history in the UK Biobank. We examined whether inferred lifetime history of depression tracked with related measures of depression and negative affect. (a) Proportion of individuals with imputed single episode (0.05%, n=791), moderate (0.10%, n=1,505), and severe (0.04%, n=631) history of depression. (b) Relative to controls, self-reported trait neuroticism was significantly greater in single-episode (*β*=0.21, p=0.045), moderate (*β*=1.93, p<2e-16), and severe depression (*β*=2.83, p<2e-16) groups. (c) Relative to controls, self-reported online depressive scores were greater among single (*β*=2.30, p<2e-16), moderate (*β*=3.19, p<2e-16), and severe (*β*=3.85, p<2e-16) depression groups. (d) Relative to controls, polygenic risk for depression was greater in single (*β*=3.7e-6, p=1.9e-9), moderate (*β*=4.7e-6, p<2e-16) and severe (*β*=5.9e-6, p<2e-16) depression groups. (e) Anti-depressant prescription rates were greater among single (*β*=0.15, p=1.5e-5), moderate (*β*=0.037, p<2e-16), and severe depression (*β*=0.24, p<2e-16), relative to controls.

**Supplemental Figure 2:**
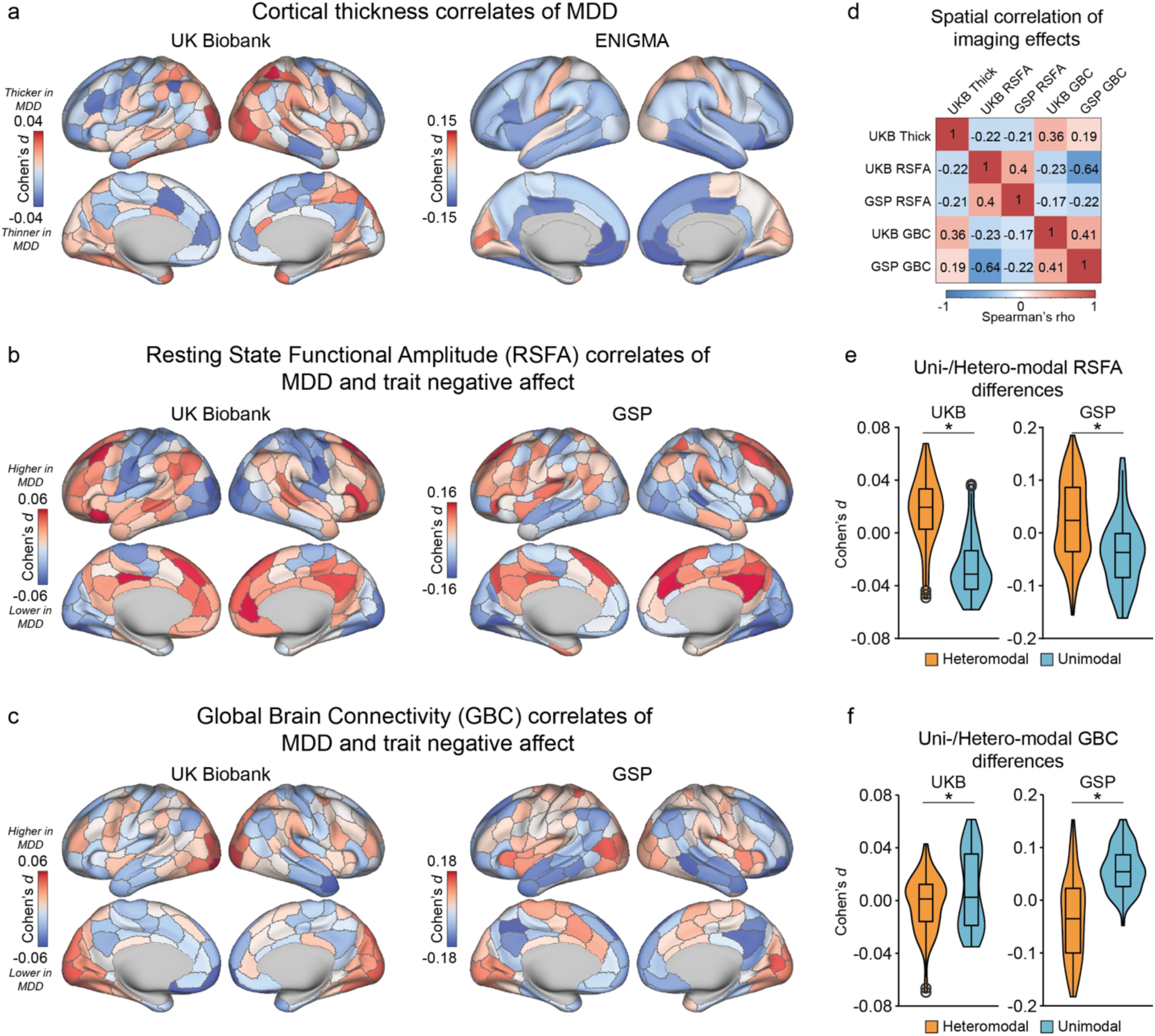
Anatomical and functional correlates of depression and trait negative affect. (a) Cortical thickness correlates of depression in the UKB (left) and ENIGMA (right). (b) Resting-state Functional Amplitude (RSFA) correlates in UKB (left) and Brain Genomics Superstruct Project (GSP; right). (c) Global Brain Connectivity correlates in UKB (left) and GSP (right). (d) Spatial correlation of depression/trait negative affect effects across modalities. Depression/trait negative affect is associated with (e) increased RSFA and (f) decreased GBC in heteromodal association cortex.

**Supplemental Figure 3:**
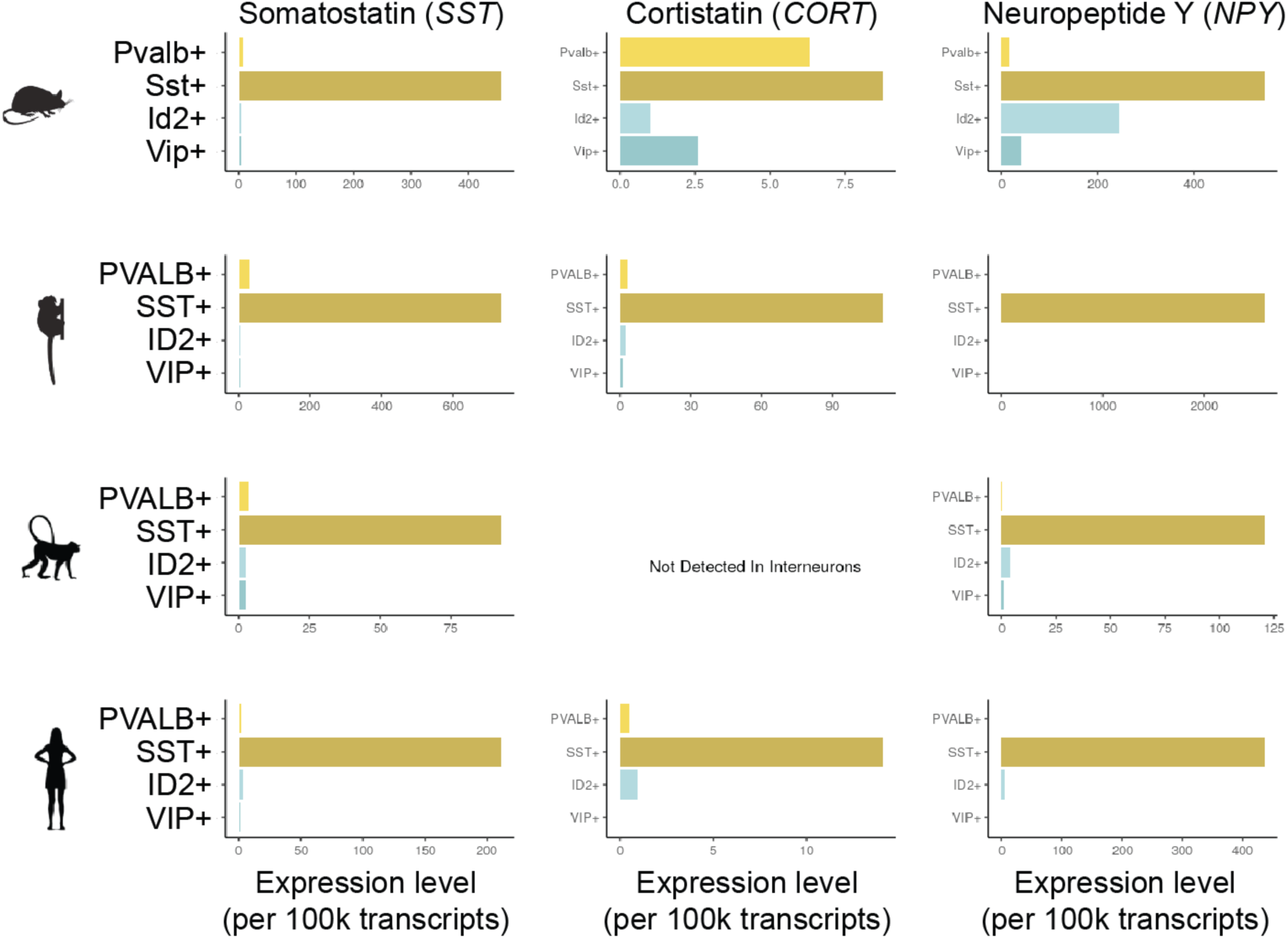
External validation of SST gene marker expression. Expression of Somotostatin (SST), Cortistatin (CORT), and Neuropeptide Y (NPY) across four interneuron populations and four species, mouse, marmoset, macaque, and human. Data are from Krienen and colleagues^105^ and can be obtained at interneuron.mccarrolllab.org/.

**Supplemental Figure 4:**
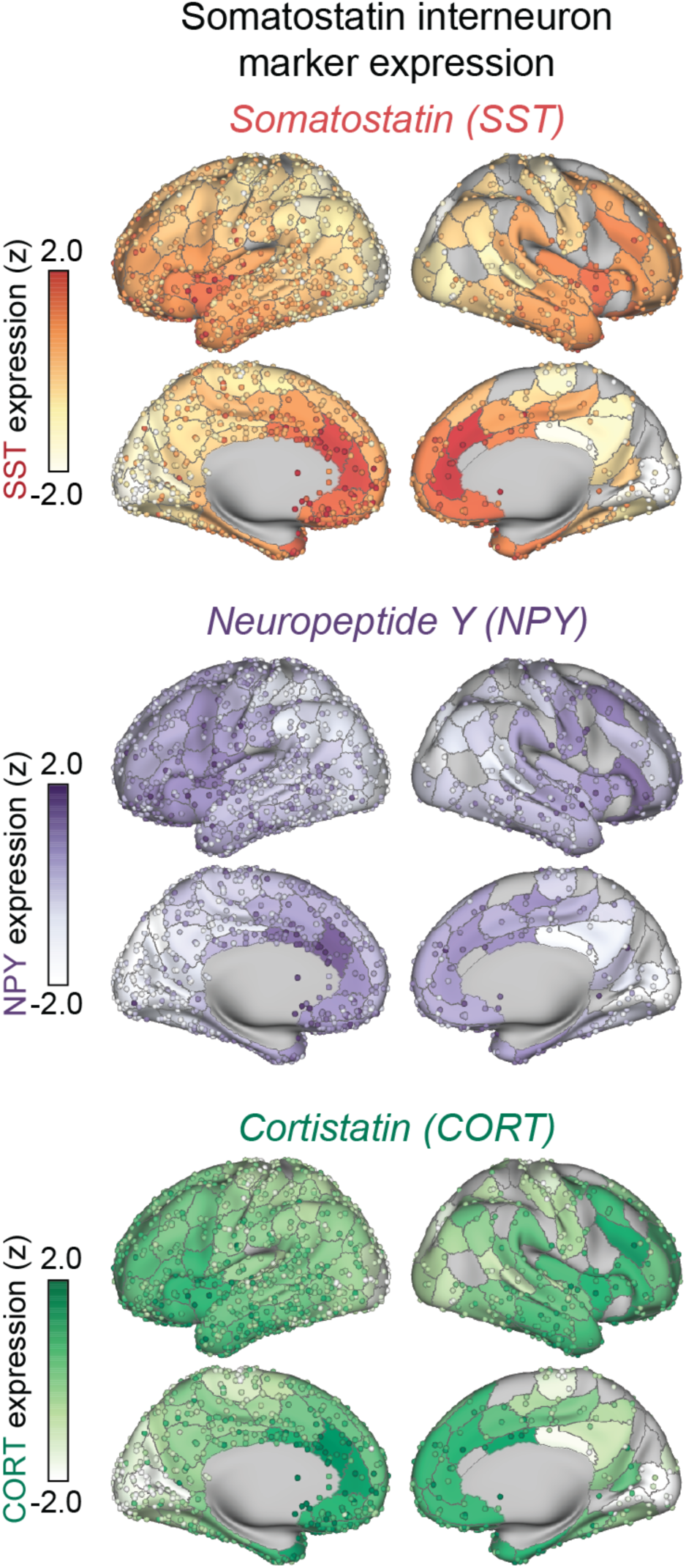
Bi-hemispheric normalized cortical AHBA expression of the SST markers Somatostatin (*SST*), Neuropeptide Y (*NPY*), and Cortistatin (*CORT*). Individual dots reflect brain tissue samples, colored by relative expression value. Parcel color reflects the average expression of individual tissue samples within a given boundary.

**Supplemental Figure 5:**
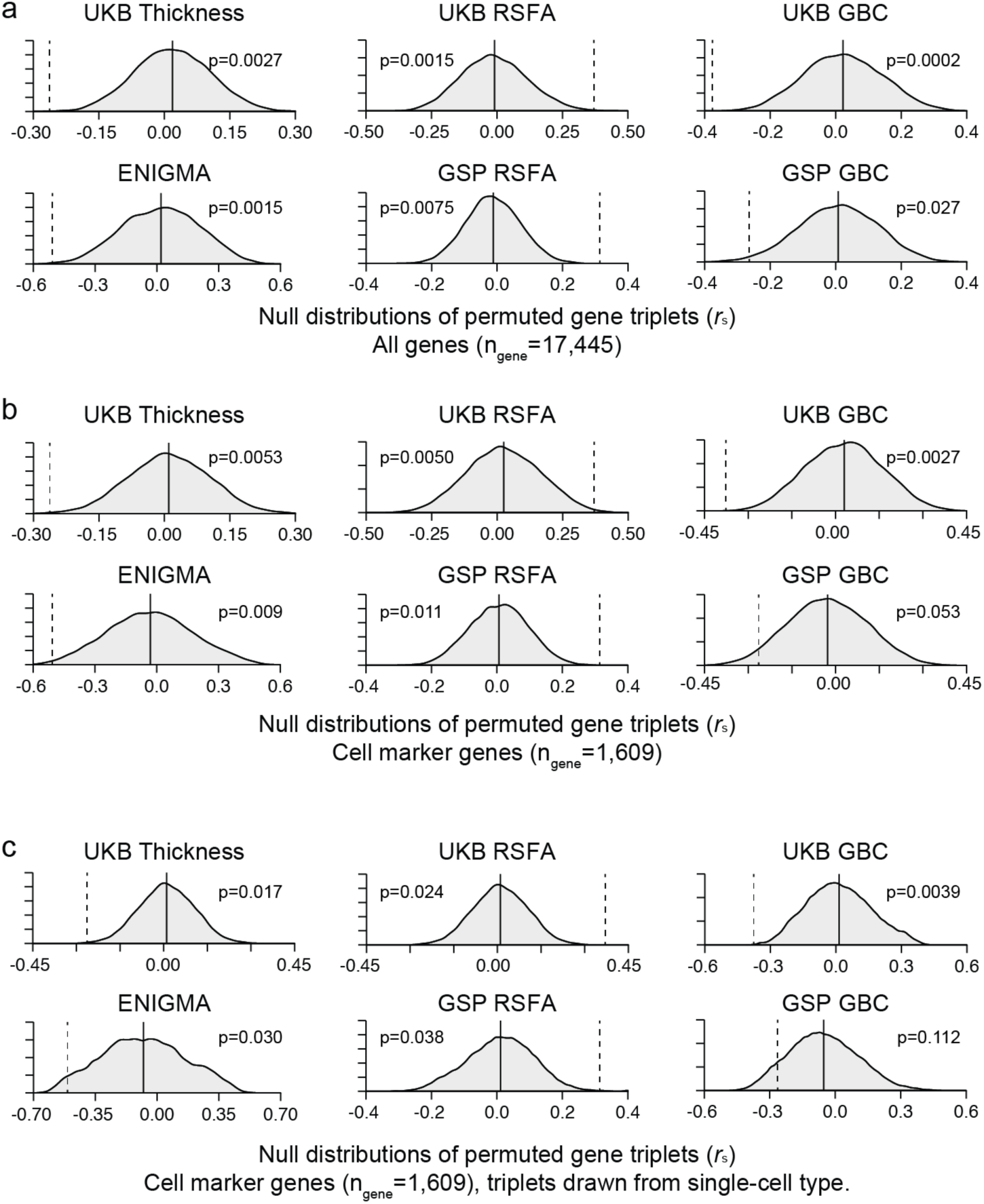
Consistent spatial correlation of SST gene markers to neuroimaging phenotypes, benchmarked against three permutation strategies. SST gene markers – *CORT*, *NPY*, *SST* – were spatially correlated to anatomical and functional correlates of depression and trait negative affect. Vertical dashed lines on reflect the strength of spatial correlation, averaged across the three SST gene markers. The strength of the SST triplet correlation was compared to three types of permutation based null distributions. (a) Null triplets were randomly selected from a pool of 17,448 genes expressed in cortex. (b) Null triplets were selected from a pool of 1,609 significant gene cell markers, defined from single-cell data from Lake and colleagues^44^ (excluding gene markers from SST cells). (c) Similarly, null triplets were selected from Lake and colleagues^44^ marker genes, however all three genes in a given null triplet were markers for the same cell type (e.g. Astrocyte, microglia, etc.)

**Supplemental Figure 6:**
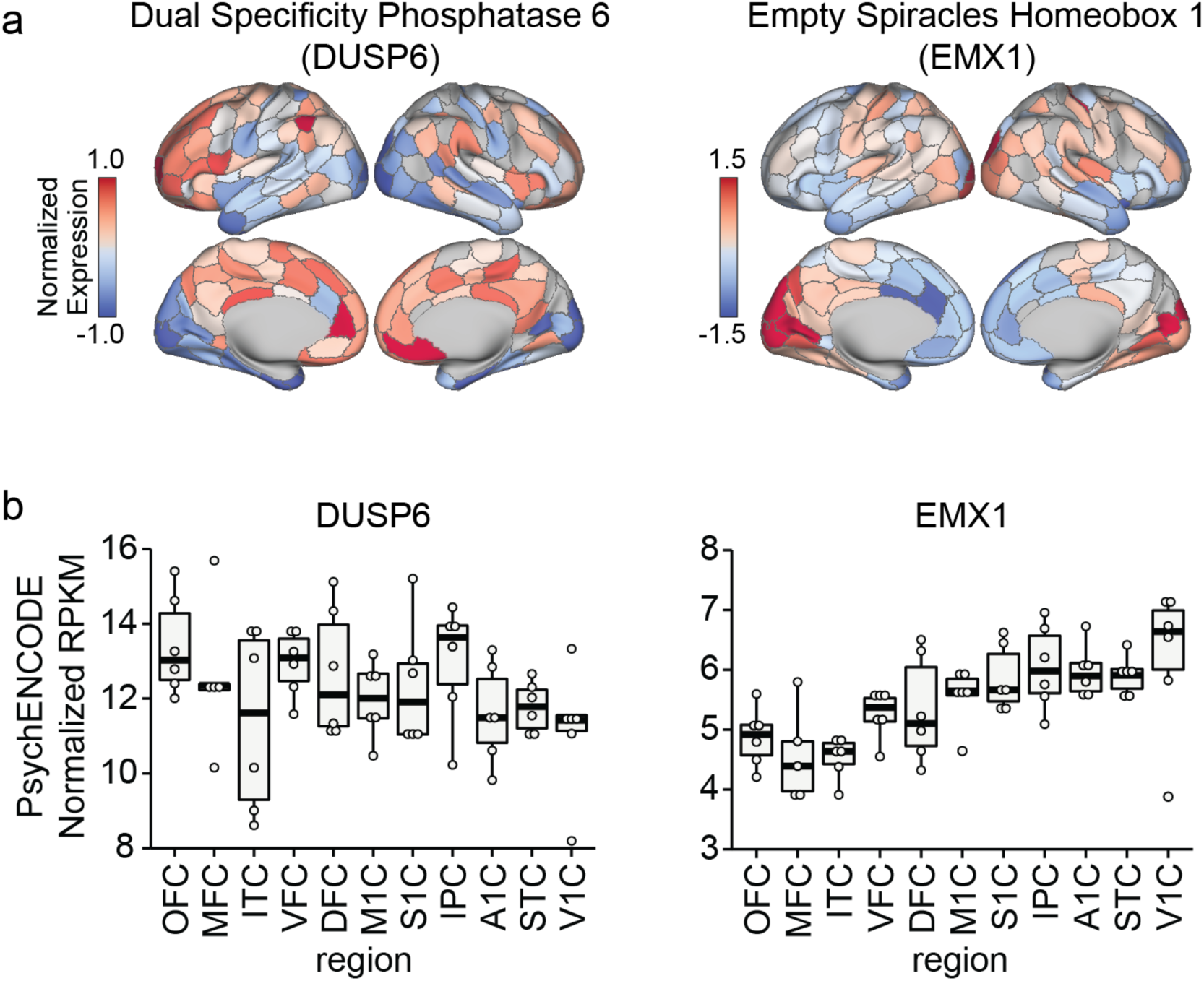
Spatial expression of *DUSP6* and *EMX1* across cortex. (a) Normalized AHBA spatial expression of *DUSP6* and *EMX1*. (b) RNAseq expression of each gene across 11 cortical regions using data from PsychENCODE^77^. RPKM values from six adult donors were batch normalized to correct for residual donor effects using ComBat^106^. Each dot represents expression in a given region, for a given donor. Regions are ordered approximately from anterior to posterior. OFC=Orbitofrontal Cortex; MFC=Medial Frontal Cortex; ITC=Inferior Temporal Cortex; VFC=Ventral Frontal Cortex; DFC=Dorsal Frontal Cortex; M1C=Primary Motor Cortex; S1C=Primary Sensory Cortex; IPC=Inferior Parietal Cortex; A1C=Primary Auditory Cortex; STC=Superior Temporal Cortex; V1C=Primary Visual Cortex.

**Supplemental Figure 7:**
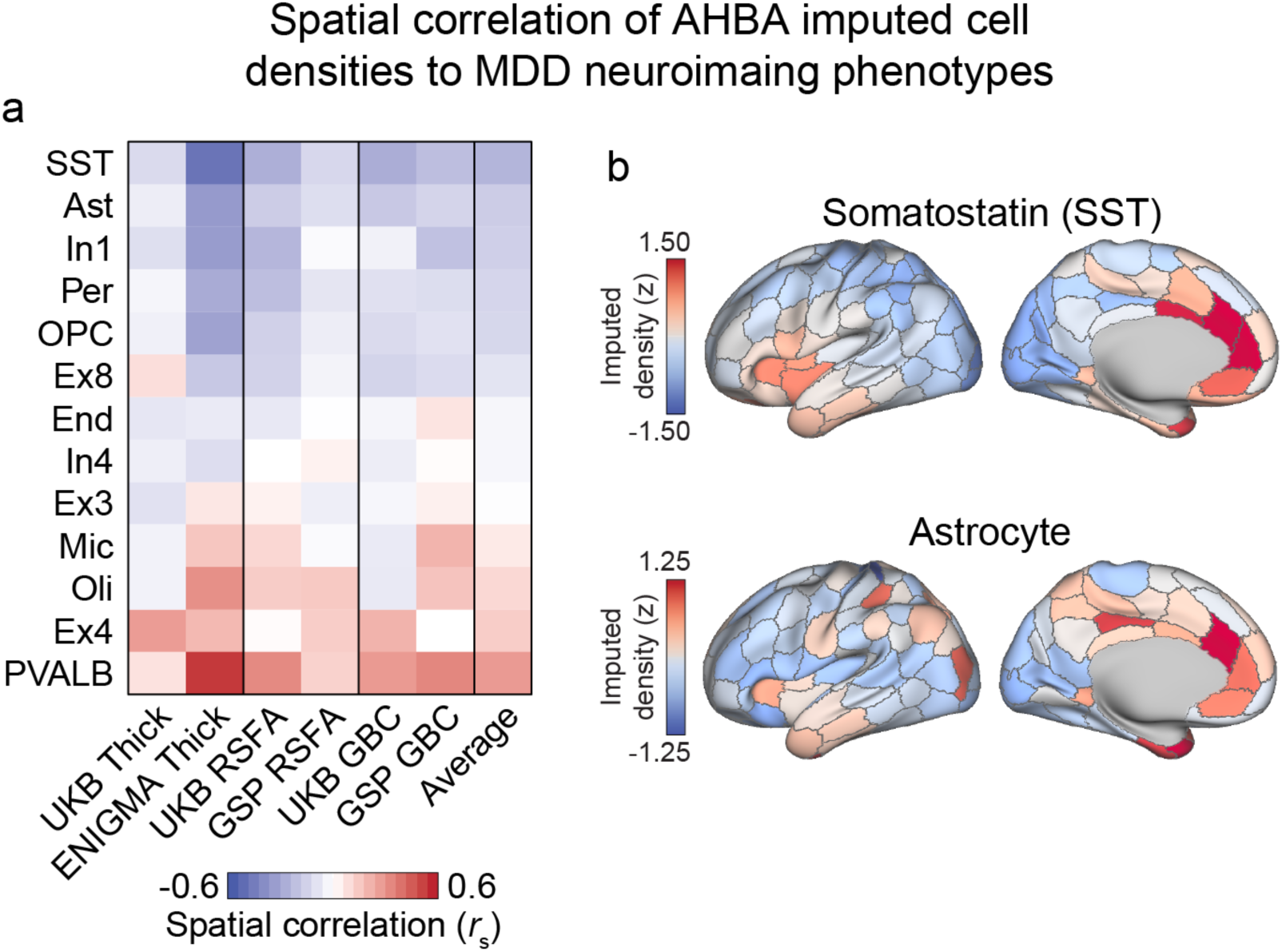
CIBERSORTx imputed cell densities correlated to depression neuroimaging phenotypes. Bulk AHBA cortical gene expression were deconvolved using single-cell data from V1C and DFC^44^. (a) Spatial correlation of each imputed density map to each dataset and modality. Negative values indicate stronger association to depression effects (e.g. increased density in areas of depression cortical thinning). Spatial density maps were imputed separately using V1C and DFC signatures before averaging across the two. (b) Somatostatin (SST) interneurons and Astrocyte imputed density maps were consistently spatially associated to the depression anatomical and functional correlates.

**Supplemental Figure 8:**
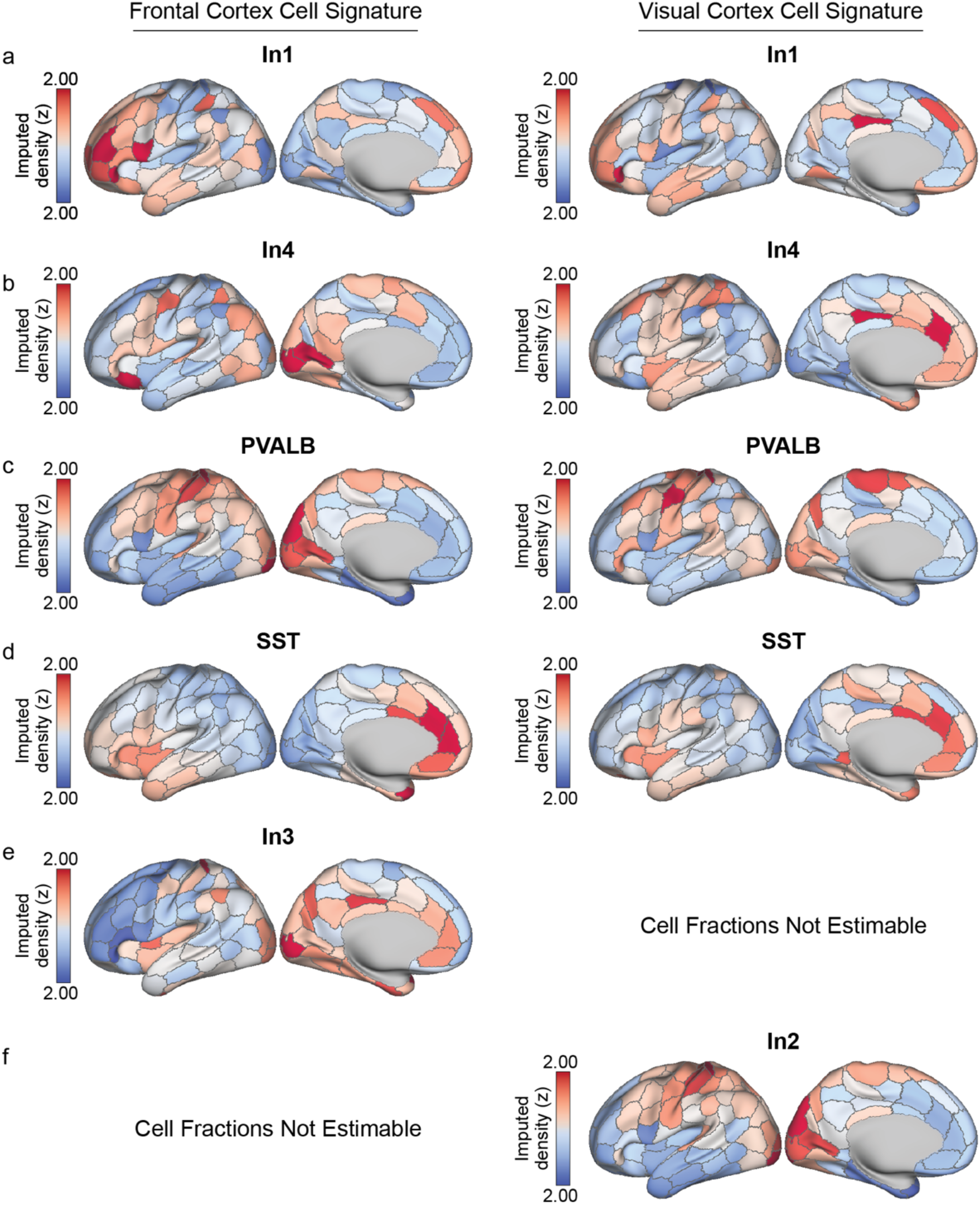
CIBERSORTx interneuron imputed spatial density maps. Imputed density maps for interneurons, estimated separately with single-cell data from V1C and DFC.

**Supplemental Figure 9:**
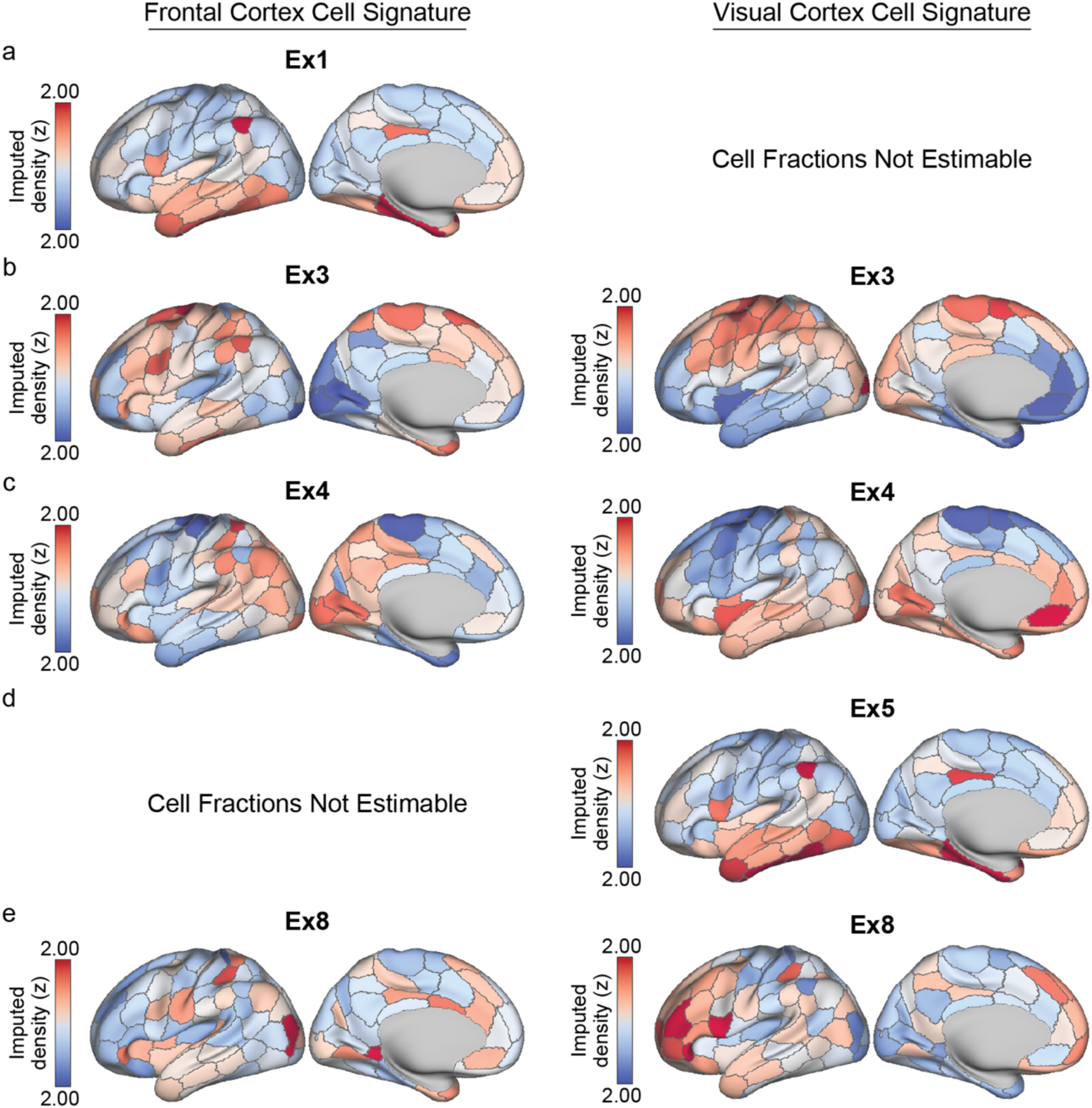
CIBERSORTx excitatory neuron imputed spatial density maps. Imputed density maps for excitatory neuron classes, estimated separately with single-cell data from V1C and DFC.

**Supplemental Figure 10:**
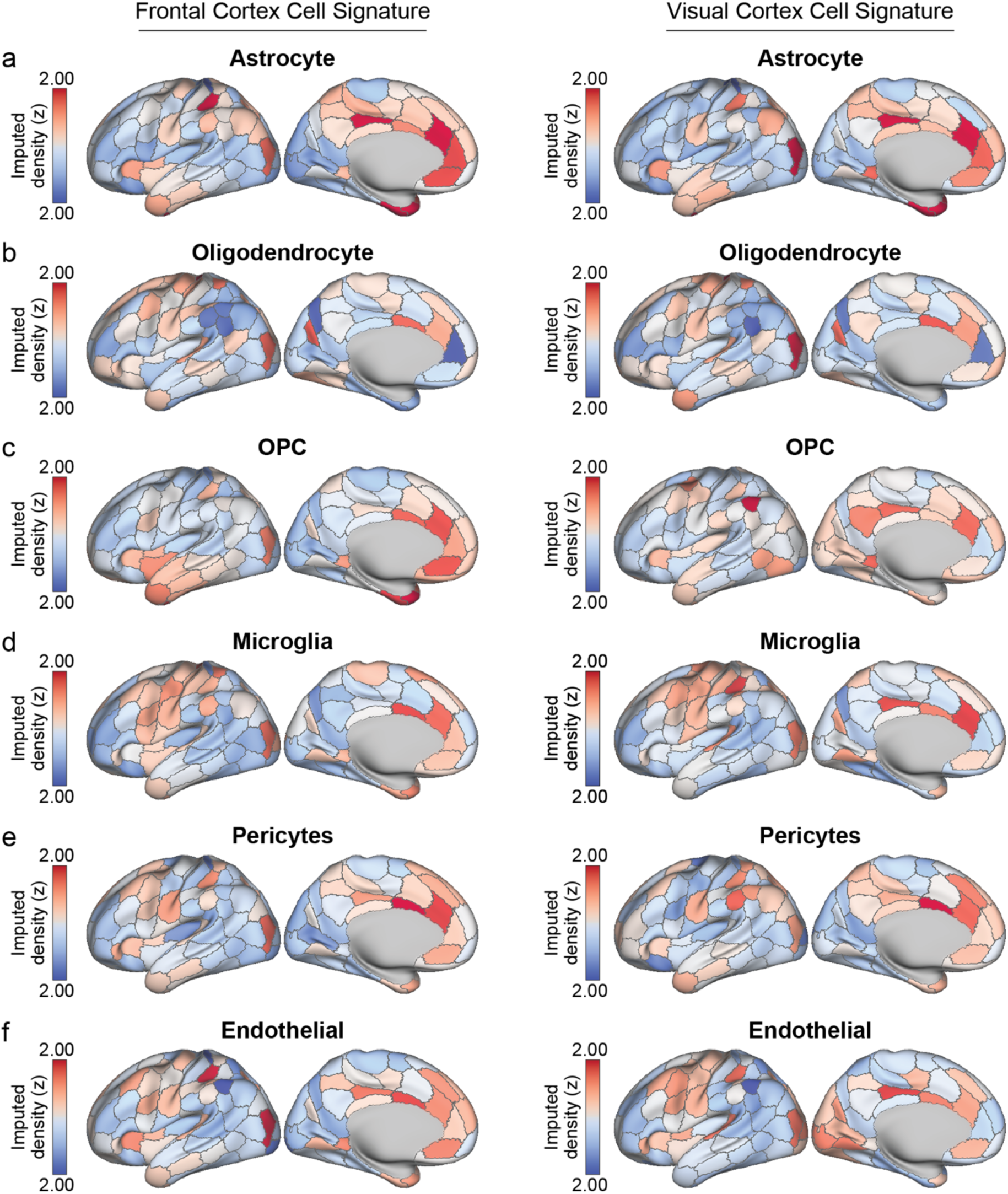
CIBERSORTx imputed spatial density maps for non-neuronal support cells. Imputed density maps for non-neuronal glia, estimated separately with single-cell data from V1C and DFC.

